# Genome-Wide CRISPR Screening Identifies Cellular Factors Controlling Nonviral Genome Editing Efficiency

**DOI:** 10.1101/2025.03.12.642795

**Authors:** Shivani Saxena, Meha Kabra, Amr Abdeen, Divya Sinha, Min Zhu, Ruosen Xie, Giovanni Hanstad, Maria A. Fernandez Zepeda, David M. Gamm, Bikash R. Pattnaik, Shaoqin Gong, Krishanu Saha

**Affiliations:** Department of Biomedical Engineering, University of Wisconsin-Madison, Madison, Wisconsin, USA; Wisconsin Institute of Discovery, University of Wisconsin-Madison, Madison, WI, USA; McPherson Eye Research Institute, University of Wisconsin-Madison, Madison, WI, USA; Waisman Center, University of Wisconsin-Madison, Madison, WI, United States; Department of Ophthalmology and Visual Sciences, University of Wisconsin-Madison, Madison, WI, USA; Department of Pediatrics, University of Wisconsin-Madison, Madison, WI, USA

**Keywords:** CRISPR, gene therapy, non-viral delivery, somatic cell genome editing, Leber congenital amaurosis, RPE, retinal cells, base editing, lipid nanoparticles

## Abstract

After administering genome editors, their efficiency is limited by a multi-step process involving cellular uptake, trafficking, and nuclear import of the vector and its payload. These processes vary widely across cell types and differ depending on the nature and structure of the vector, whether it is a lipid nanoparticle or a different synthetic material. We developed a novel genome-wide CRISPR screening strategy to better understand these limitations within human cells to identify genes modulating cellular uptake, payload delivery, and gene editing efficiency. Our screen interrogates the cellular processes controlling genome editing by Cas-based nuclease and base editing strategies in human cells. We designed a genome-wide screen targeting 19,114 genes in HEK293 cells, and we identified six genes whose knockout increased nonviral editing efficiency in human cells by up to five-fold. Further validation through arrayed knockouts of the top hits from our screen boosted the editing efficiency from 5% to 50% when Cas9 was delivered via lipid-based nanoparticles. By designing the guides to target the screen library cassette, we could accurately track the library sgRNA identity and the editing outcome on the same amplicon via short-read sequencing, enabling the identification of rare outcomes via ‘computationally’ sorting edited from unedited cells within a heterogenous pool of >200M cells. In patient-derived human retinal pigment epithelium cells derived from pluripotent stem cells, BET1L, GJB2, and MS4A13 gene knockouts increased targeted genome editing by over five-fold. We anticipate that this high-throughput screening approach will facilitate the systematic engineering of novel nonviral genome editing delivery methods, where the identified novel gene hits can be further used to increase editing efficiency for other therapeutically relevant cell types.

## Introduction

Therapeutic genome editing has entered clinical trials^1–3^, with candidates progressing to late-stage clinical trials^4^ and one reaching approval^5,6^. Many delivery vehicles, such as lipid nanoparticles^7–9^, virus-like particles^10,11^, and viral vectors^12–14^, can be loaded with CRISPR components, enabling the development^3^ of new genome editing tools, such as base editors within lipid nanoparticles^15,16^. While these interventions could reduce suffering for millions of Americans affected by disease^17,18^, a key challenge for the field is fundamentally understanding how these approaches generate both intended and unintended genomic outcomes. Editing somatic cells ex vivo^19^ has proven efficient (>50%) with these vehicles, leading to the first approved CRISPR therapy, Casgevy^20^. However, achieving gene editing in specific cell types within tissues, such as sensitive sensory post-mitotic cells in the inner ear^21^ or eye^22^, remains challenging. For non-viral vehicles ^23^, efficiency typically hovers around 25%. Effective phenotype rescue in post-mitotic cells, like photoreceptor cells, generally require editing efficiencies of around 50% or higher to ensure a substantial therapeutic effect^24^. Therefore, optimizing non-viral delivery methods to achieve higher editing rates in post-mitotic cells and precise genome editing in patient tissues is crucial.

When the delivery vector reaches the cell membrane, various factors can influence the route taken by the vector to cross the cell membrane, which also may affect the vector’s potency to deliver the genome editor (**Figure 1A**). Prior work^25,26^ investigating endocytosis pathways and endosomal escape for various small molecules such as siRNA and CRISPR-Cas9^27^ indicated that less than 1% of the cargo made it to the cell’s cytosol. Limitations in our understanding of delivery are partly due to technical challenges associated with profiling the signal transduction pathways and nuclear import of the delivery vehicle. Many studies have reported on non-viral strategies elucidating Cas9 Ribonucleoprotein (RNP) internalization and its biodistribution across tissue^7,28,29^. However, such efforts typically do not assay both editing efficiency and delivery process within the same cell.

**Figure 1:**
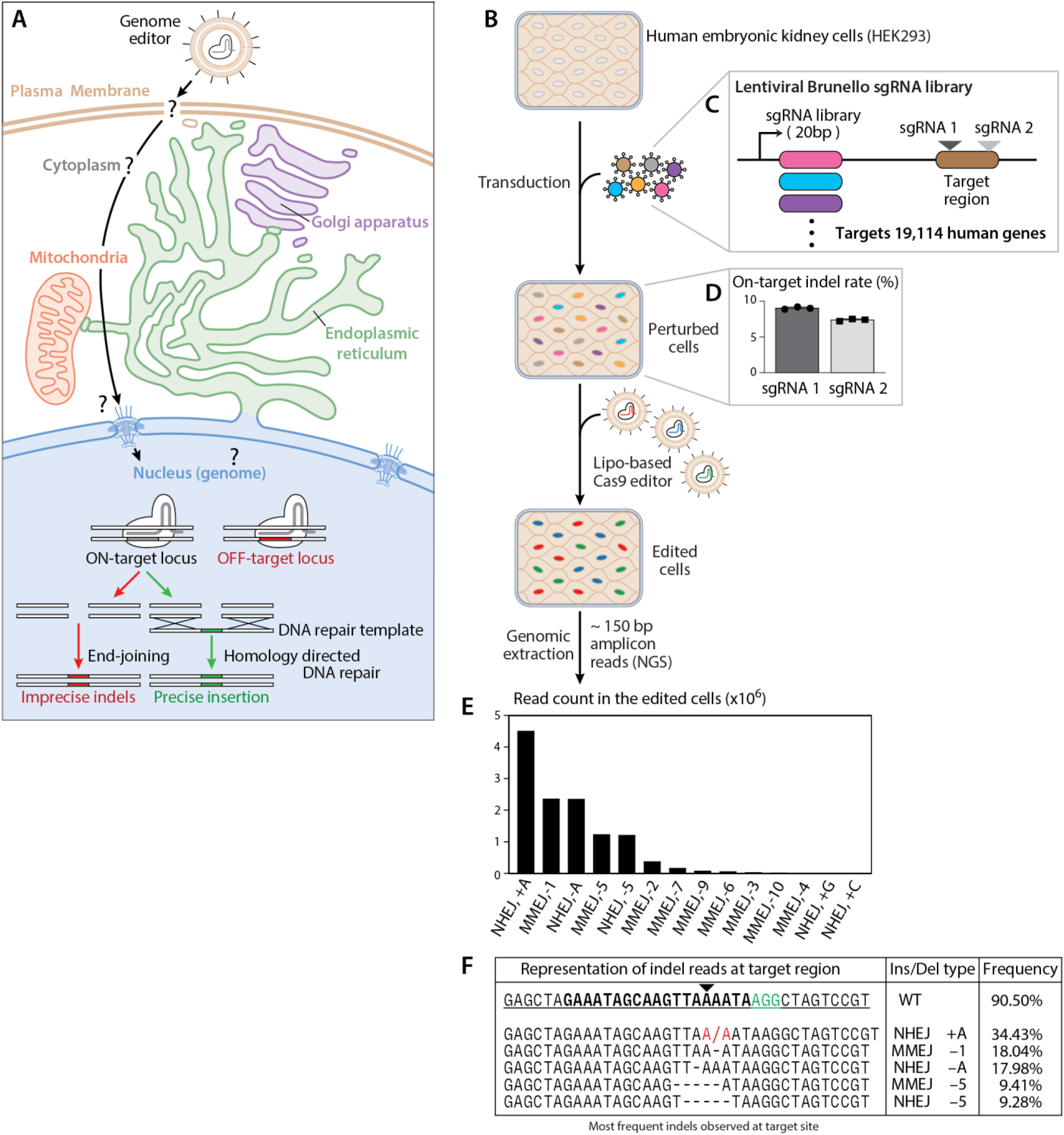
Genome-wide CRISPR screening to identify genes that increase nonviral genome editing. **(A)** Several types of editing outcomes in therapeutic genome editing. **(B)** Design of the screen workflow. Lentivirus transduction of HEK293 cells integrates the Brunello transgene expressing *Sp*Cas9 protein and sgRNA under EF1-alpha promoter. A total of ∼200 million cells were transduced and subjected to puromycin selection. This transduced Brunello-HEK cell pool was subjected to nonviral editing by lipofectamine 2000 with a *Sp*Cas9-sgRNA ribonucleoprotein complex targeting the blue target region in the Brunello transgene. **(C)** Schematic of LentiCRISPRv2-Brunello vector expressing one sgRNA within a genome-wide library (targeting 19,114 genes; 4 sgRNAs per gene) and *Sp*Cas9 nuclease. The downstream protospacer and protospacer-adjacent motif (PAM) region within the Brunello cassette is highlighted as the target for nonviral editors. **(D)** CRISPR based-editing efficiencies measured via deep sequencing are shown as the percentage of indel reads (y-axis) for the two guides (sgRNA1 and sgRNA2, **Table S1**) used to target the internalized Brunello cassette in HEK293 cells (n=3). **(E)** The indel pattern shown here for different types of repair mechanisms — MMEJ (±) and NHEJ (±) is plotted against the total read count in the edited group from the screen cell population. Representation of indel reads from the edited group with NHEJ (+A) being the most prevalent indel pattern induced via both guide RNAs. **(F)** Representation of indel profile at the target region (sgRNA in bold letters with a PAM sequence “AGG” in green) of the internalized Brunello cassette, showing the percentage frequency of the type of indel outcome, highlighted in red. RNP, Ribonucleoprotein; UTF, Untransfected; KO, Knock-out; NHEJ, Non-Homologous End-Joining; MMEJ, Microhomology Mediated End-Joining.

Apart from delivery, CRISPR-Cas9 employs a guide RNA to direct the Cas9 nuclease to a precise genomic location, where the Cas9 nuclease cleaves the DNA, inducing a double-strand break (DSB). In human cells, these DSBs are then primarily repaired either through non-homologous end joining (NHEJ)^30,31^, which can introduce insertions or deletions or via homology-directed repair (HDR)^31^, which can incorporate a template DNA sequence for precise edits (**Figure 1A**). Strategies to improve efficiency can involve modulating DNA repair pathways and engineering guide RNAs to enhance editing precision and reduce off-target impacts, making nuclease-based editors crucial yet complex tools in genome engineering. For example, adding the 53BP1 inhibitors^32^, DNA-PKcs inhibitors^33–35^, and RPA70 dsDNA binding inhibitor^33^ to the culture media have enhanced the efficiency of CRISPR-mediated editing and modulated the DNA DSB repair pathways^36,37^ in human cell types, including embryonic stem cells. Genes controlling DNA repair include DNA damage response genes, such as *POLQ*^36,38^, *RAD18*^37,39^, *MRE11*^36,40^, and *LIG3*^41,42^.

Pooled genome wide CRISPR screens facilitate the systematic mapping^21,34,41–45^ of gene determinants in a biological process. The CRISPR screens perturb a gene by knockout, activation, and repression using large libraries of sgRNAs designed to target a specific subset of genes within the genome. We reasoned that pooled CRISPR screening using a sgRNA library targeting every gene in the human genome could uncover genetic factors that influence nonviral gene editing in human cells, specifically those affecting CRISPR Cas9 editing efficiency. Genes can be identified that influence editor trafficking and the genome editing processes, connecting gene functions and their biological phenotypes in a single experiment.

Here, we conducted a genome-wide pooled genetic screen to identify 26 genes among 19,114 human genes that — when knocked out — could increase the frequency of on-target gene edits by Cas9. Across several different human cell types, the knockout of each of the 26 genes could increase the editing of Cas9-nuclease and Cas9-base editors packaged in several nonviral vehicles delivering mRNA or protein. Importantly, strategies targeting these identified genes could enhance CRISPR editing efficiency in clinically relevant post-mitotic cells, potentially enabling mutation correction in patient tissues. These findings will likely contribute to developing potent nonviral genome editing formulations suitable for *in vivo* applications.

## Results

### Design of genome-wide pooled CRISPR screen

CRISPR screening studies have been used to investigate DNA repair profiles and uncover new functions for canonical DNA damage response^37,43^ and repair genes^32,36,44–46^. We reasoned that a broader screen could uncover genetic perturbations that change the uptake and intracellular trafficking of CRISPR delivery vehicles and payloads within the cells. Inspired by these and other studies^32,47–49^, we designed a pooled CRISPR screening method to interrogate the effects of nearly all human genes (∼20,000 genes) on the editing efficiency of CRISPR-Cas9 nucleases using nonviral delivery. For this high-throughput screening approach, we employed a well-characterized LentiCRISPRv2 vector with a Brunello sgRNA library^53^, which targets 19,114 human genes with four sgRNAs per gene^54^. We designed an sgRNA targeting the internalized LentiCRISPRv2-Brunello cassette integrated within the HEK293 cells via lentivirus transduction using a low multiplicity of infection (MOI ∼ 0.3) (**Figure 1B**). We delivered CRISPR Cas9 RNP complexed with the sgRNA, targeting the LentiCRISPRv2-Brunello cassette, producing indels proximal to the Brunello sgRNA sequence (∼20 bp, perturbation identifier). Successful nonviral editing within the HEK293-Brunello perturbed cell population would produce indels at the target region within the LentiCRISPRv2-Brunello cassette. Nonproductive delivery or editing would leave the target region untouched. Altogether, we can determine the editing outcome of each allele within the perturbed pool of cells (∼200 million cells) by extracting genomic DNA and sequencing the ∼ 150 bp Brunello transgene via next-generation sequencing. Paired-end, short-read amplicon sequencing of this PCR amplicon (∼150 bp, Brunello transgene) containing the indel outcome and the perturbation identifier (guide library sgRNA) provides base pair resolution for many alleles within the experimental pool of cells. Importantly, the sequenced read includes both the editing outcome, i.e., NHEJ induced indel sequence and the sgRNA used to knock out the host cellular determinant (19,114 Brunello library genes), within the same amplicon. The enrichment of the perturbation identifier (Brunello sgRNA), associated with successful NHEJ-mediated indel formation in the pool of non-virally edited cells, ensures a high level of confidence in its role to influence NHEJ-mediated editing.

We evaluated this strategy within human embryonic kidney (HEK293) cells transduced at a low MOI (∼0.3) with the Brunello vector library^50^, comprising 76,456 targeting sgRNAs, along with 1000 non-targeting control sgRNAs. After selecting transduced cells, we added *Sp*Cas9 RNPs delivered via lipofectamine to produce edits at the target region within the LentiCRISPRv2-Brunello cassette (**Figure 1C**). Four days after the addition of lipofectamine, to allow for indel formation by Cas9 RNPs, genomic DNA was isolated from the pool of cells (See Methods). We observed 16.4% ± 2% editing (edited group, via both guide RNAs) in the total pool of cells targeting the Brunello cassette (**Figure 1D**). PCR amplification and short-read sequencing revealed a diverse distribution of indels on the target region of the cassette (**Figure 1E**). In our genome-wide screen, we mapped 17,988 genes from the Brunello library, with each sgRNA having a median of ∼200 reads covered per gene (**Supplementary Figure S1A**). Additionally, sequencing identified 1,126 genes that were depleted from the total screen population.

By analyzing the reads, NHEJ-induced indels were highly reproducible. They exhibited similar editing patterns targeted by two distinct sgRNAs with their protospacers and PAMs located within the same integrated Brunello transgene (**Figure 1F**). We then segregated the read data into two distinct allele sets based on outcomes: one identified by NHEJ-induced indels (NHEJ +), labeled as the “edited allele set,” and another set without any indels at the on-target site (NHEJ -), designated as the “control unedited allele set.” We then used the MAGeCK^51,52^ algorithm (Model-based Analysis of Genome-wide CRISPR/Cas9 Knockout) to quantify a ‘phenotype’ for each sgRNA in our library. The phenotype is defined as fold-change enrichment of a particular sgRNA in the edited set of reads relative to the control unedited set of reads. sgRNAs with positive fold-change phenotype indicates that gene activity suppresses editing, while negative fold-change phenotype indicates that gene activity promotes editing. Sequencing the amplicons deeply to obtain a high coverage enabled statistically meaningful comparisons between the edited group and the control group (**Figure 2A**). These two groups were generated computationally by extracting edited reads from the total reads sequenced, leaving the unedited reads designated as the control read group.

**Figure 2:**
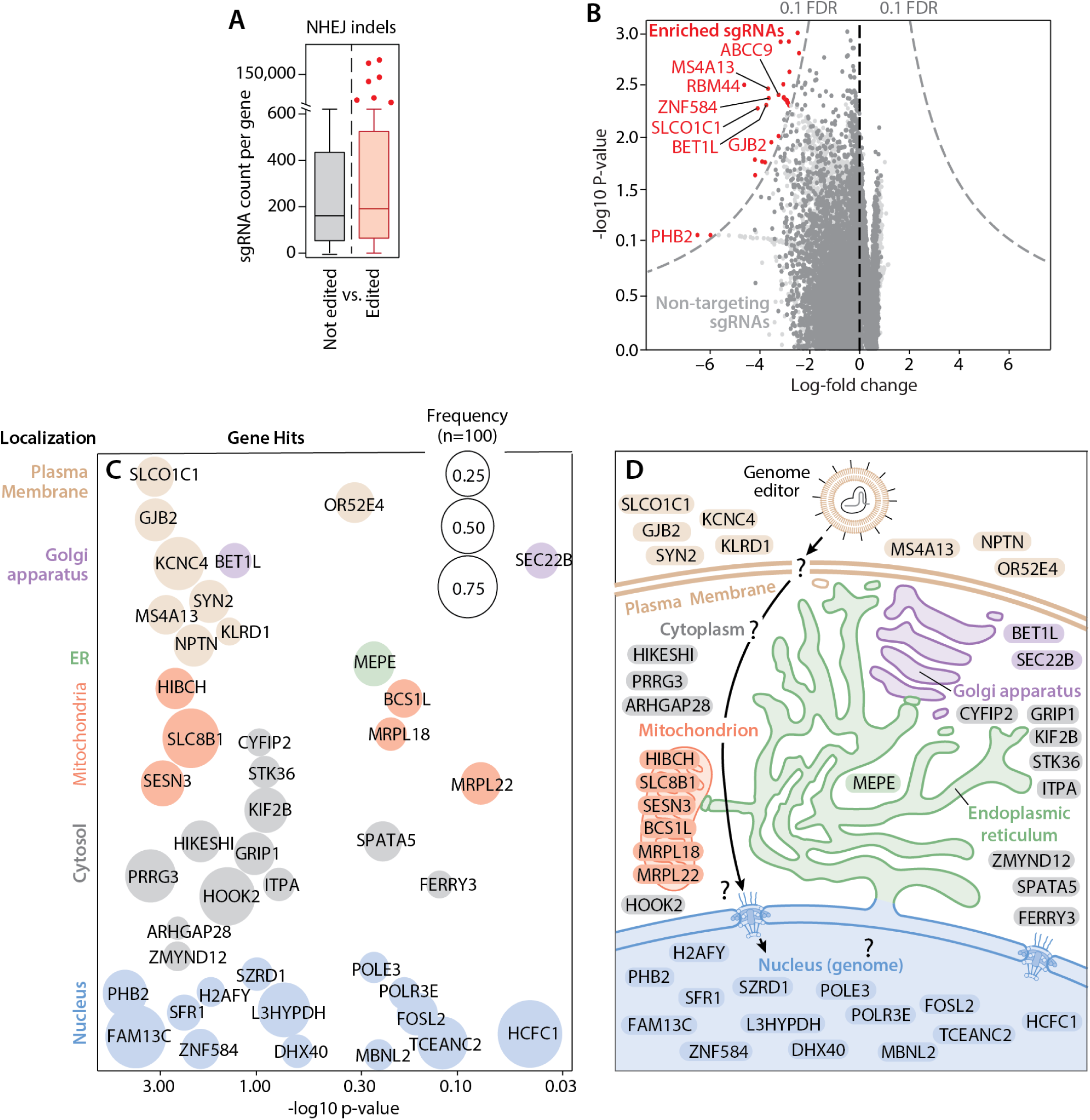
Analysis of the genome-wide pooled screen. See also Figure S1. **(A)** Total sgRNA coverage (identified from the genome-wide screen) is shown in both groups analyzed via MAGeCK analysis (n=100), between the control group (subsample) and the edited group, reporting a median of ∼ 200 sgRNA reads per gene. **(B)** Volcano plot showing the significance and log-fold change for Brunello sgRNA gene targets in cells with indels at the on-target site. Top sgRNAs (corresponding to respective genes) enriched in the screen were selected from the screen due to their role in suppressing NHEJ in HEK293 cells, and depletion of these sgRNAs in promoting NHEJ. The scatterplot shows log-fold change (x-axis) against the log-P-values (y-axis) for specific genes, where the enriched genes are shown in red and non-targeting genes are shown in grey. **(C)** The bubble plot represents the multiple iterations done via the MAGeCK algorithm (n=100), where top gene hits on the screen are plotted against their p-value. The size of the bubble corresponds to the frequency of occurrence for that particular gene and color represents the cluster defining the subcellular localization of the genes− grey, Cytosol; green, ER; purple, Golgi Apparatus; red, Mitochondria; blue, Nucleus; brown, Plasma Membrane. **(D)** Schematic of the genome editing cargo trafficking into the nucleus of the cell, affected by the different host genes uncovered in the screen. RNP, Ribonucleoprotein; UTF, Untransfected; KO, Knock-out; NHEJ, Non-Homologous End-Joining; MAGeCK, Model-based Analysis of Genome-wide CRISPR/Cas9 Knockout: FDR, False Discovery Rate.

To address the sample size disparity between the two sets, we divided the control set into random subsets (similar size as the edited allele set) and conducted multiple iterations (n=100) of the MAGeCK analysis (**Supplementary Figure S1B**). Comparing the effects of all gene targets within the Brunello library, among the two subsets (edited and unedited reads), revealed genes significantly enriched in the edited allele set, shown as hits (**Figure 2B**). At the genomic level, genes corresponding to the respective sgRNA that are consistently enriched in the edited group across multiple iterations (n=100) are represented by ‘frequency,’ defined as the total number of times a gene was identified as significant. The frequency of these gene hits, grouped by their subcellular localization, is plotted against their significance, as illustrated in **Figure 2C**. We curated a list of 26 genes based on their distinct functions (**Supplementary Table S1**), clustered based on the subcellular localization of the genes and their frequency of occurrence in the multiple MAGeCK analysis iterations. The central hypothesis of our screen was to explore novel assemblies of CRISPR-Cas9 particles influenced by the activation or suppression of various genes (**Figure 2D**). This approach enables the investigation of distinct biological processes, including cellular trafficking, DNA nicking, DNA double-strand break formation, and DNA repair mechanisms involved in genome editing in human cells.

### Validation studies identified six top genes exhibiting increased on-target editing

We evaluated 26 hits from a compiled roster featuring the 43 most potent sgRNAs, all implicated in DNA repair and intracellular trafficking mechanisms. To assess the hits’ effects at a genomic level, we performed an arrayed CRISPR screen in the HEK293 cell line. We cloned each sgRNA for the hit gene into a separate LentiCRISPRv2 vector and established polyclonal monogenic KO lines using HEK293 cells (**Figure 3A**). Next, we repeated lipofectamine nonviral delivery with these polyclonal KO lines using a validated sgRNA and *Sp*Cas9 RNP, inducing a DSB within an endogenous locus. To evaluate the NHEJ indel outcomes in the 26 mutant lines (KO-HEK293 cells) at the genomic level, we compared them with NHEJ editing in WT-HEK293 cells in a single arrayed screen experiment. We targeted the *NRL* gene, which encodes for the neural retina-specific leucine zipper protein in human cells, and induced NHEJ via CRISPR-Cas9. Lipofection was used to introduce NHEJ via *NRL* sgRNA (**Figure 3B**) simultaneously at a specified locus across 26 mutant lines. The editing rate in WT-HEK293 cells served as a baseline to compare the NHEJ editing efficiency in all the KO-HEK293 lines. We then performed next-generation sequencing to quantify the indel outcomes at the *NRL* locus, observing an increase in editing for some of the mutant lines (**Figure 3C**). We observed at least six mutant HEK293 lines with a ∼2-fold increase in NHEJ editing at the endogenous *NRL* locus. The editing rates were intentionally maintained at low levels to assess the extent of potential increases in editing efficiency.

**Figure 3:**
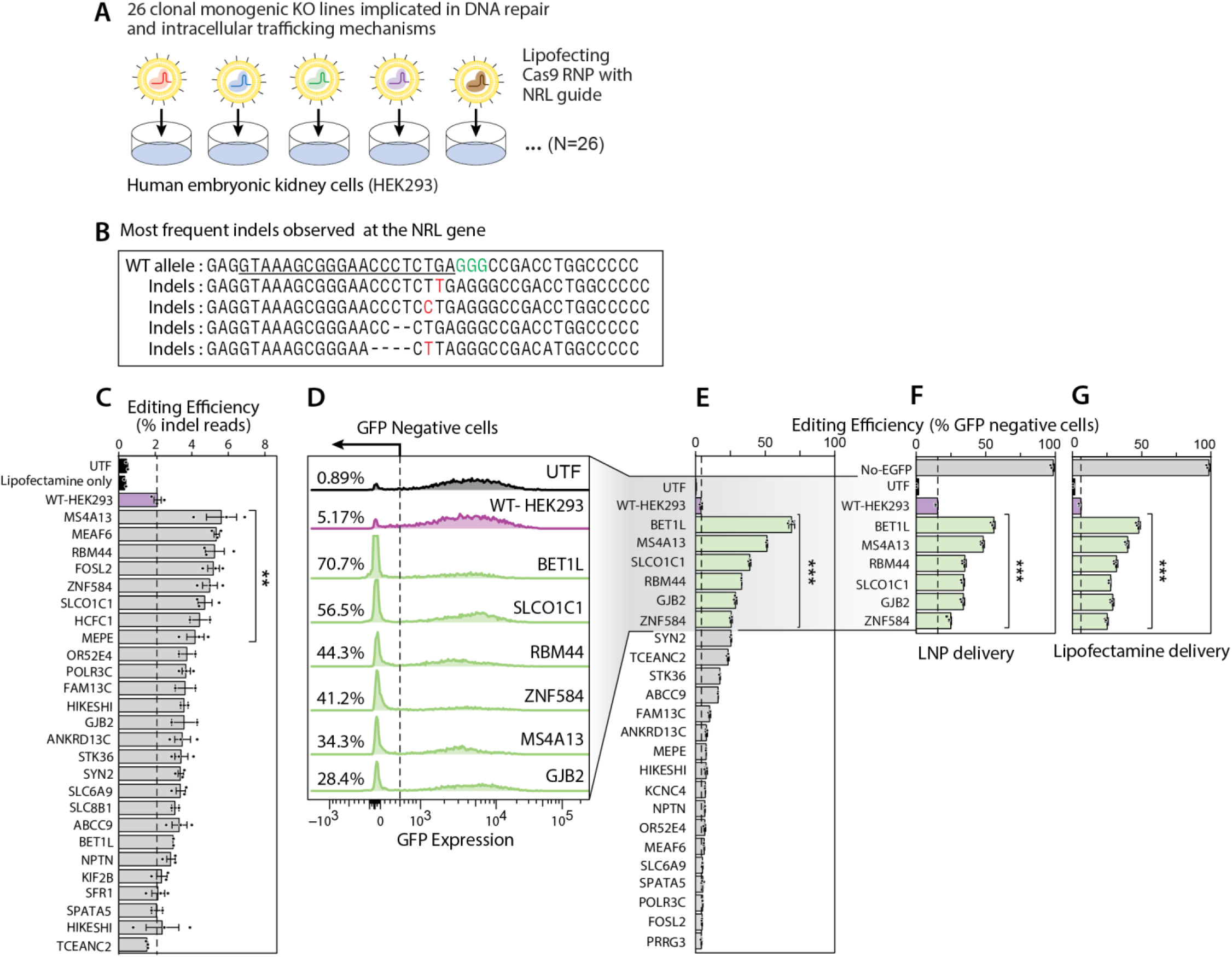
Arrayed screen for validating hits as potent enhancers of CRISPR-mediated NHEJ. See also Figure S2. **(A)** The schematic illustrates the experimental workflow for generating 26 monogenic knockout (KO) lines in HEK293-GFP cells. The LentiCRISPRv2 plasmid was used for knocking out 26 different ‘gene hits’, using a KO cassette expressing *Sp*Cas9 and the guide RNA for the target gene. The plasmid also conferred puromycin resistance, which was used to select edited cells post transfection, for the validation screen. The established KO lines were further edited using lipid-based methods (lipofection/LNPs) to target an endogenous locus at the NRL gene. **(B)** Representation of the indel reads at the endogenous locus at NRL gene post-editing, measured via next-generation sequencing. **(C)** On a genomic level, the 26 different monogenic KO lines were edited using CRISPR Cas9 RNP, targeting the *NRL* gene locus. Data is shown for three replicates per KO HEK293 line (n=3). **(D)** Representative histogram plot shows the knockout of GFP protein expression 96 hours post-treatment with LNPs. The plot displays top genes significantly increasing the knockdown of GFP protein: *BET1L*, *MS4A13*, *RBM44*, *SLCO1C1*, *GJB2*, and *ZNF584*. The X-axis represents GFP protein expression levels, while the Y-axis represents the percentage of cells negative for GFP expression. UTF samples are shown as a black histogram (negative control), and WT-HEK293 samples are shown as a pink histogram (positive control). **(E)** GFP expression knockdown is shown for 23 genes as the percentage of cells negative for GFP protein (Y-axis). Data is presented as an average of three replicates for each monogenic KO HEK293-GFP line (n=3). UTF samples are represented by black bars, and WT-HEK293 (positive control) samples are represented by pink bars in all plots. **(F)** HEK293-GFP cells knocked out for the top six genes validated in the arrayed screen were edited using lipid nanoparticles (LNP). Editing was performed by delivering CRISPR Cas9 mRNA and GFP KO guide for six KO lines. Data is shown for three replicates (n=3) per condition (six KO lines, WT-HEK293 cells, and UTF). **(G)** HEK293-GFP cells knocked out for the top six genes validated in the arrayed screen were edited via lipofection. Editing was performed by delivering CRISPR Cas9 RNP and GFP KO guide for six KO lines. Data is shown for three replicates (n=3) per condition (six KO lines, WT-HEK293 cells, and UTF). Statistical significance was calculated using ordinary one-way ANOVA. Dunnett’s multiple comparison test was used as the post-test (C, D, E, and F). *, p < 0.05; **, p < 0.01; ***, p < 0.001; ****, p < 0.0001. ANOVA, Analysis of Variance. RNP, Ribonucleoprotein; UTF, Untransfected; KO, Knock-out; NHEJ, Non-Homologous End-Joining.

Furthermore, we diversified the payload and vehicle, by evaluating mRNA-based sgRNA delivery via lipid nanoparticles targeting EGFP in a fluorescent reporter HEK cell line, which constitutively expresses a single transgene copy of the EGFP protein. A flow-cytometry-based assay was designed for the ease of testing multiple screen hits and evaluating the editing at a protein level. To investigate enhanced NHEJ, we compared GFP protein knockdown rates following lipid-based nanoparticle (LNP) delivery of Cas9 mRNA and EGFP sgRNA across 23 mutant lines, including positive and negative controls. This allowed us to establish a baseline indel frequency for comparison with the mutant lines (**Supplementary Figure S2**). We analyzed the percentage of GFP-negative cells using flow cytometry to quantify editing efficiency at the protein level. Interestingly, we identified six genes—*BET1L, MS4A13, RBM44, SLCO1C1, GJB2, and ZNF584*—that exhibited more than a tenfold increase in GFP knockdown in HEK293 cells, compared to the expression in WT-HEK293 cells (**Figures 3D and 3E**). We saw a similar trend of increased editing for these six genes, which remained consistent at the protein and genomic level.

Furthermore, we evaluated two lipid-based non-viral delivery (LNP; **Figure 3F** and Lipofectamine; **Figure 3G**) strategies to deliver the sgRNA and editor payload, successfully verifying the reproducibility across delivery platforms for the top gene hits identified in this validation screen. We concluded that indel outcomes generated by validating common genes across two delivery platforms produced enhanced editing, which informed a reproducible editing strategy in harder-to-edit cells. In conclusion, we selected the six genes consistently exhibiting increased editing across both delivery strategies.

### Knocking out BET1L and MS4A13 enhanced the nuclear localization of the Cas9-GFP protein

To characterize the effects of three genes – *BET1L, MS4A13*, and *GJB2*, it was imperative to investigate the intracellular trafficking of the editor protein post-delivery. After observing no change in editing via electroporation (**Supplementary Figure S1C**) to knock down GFP expression in the mutant HEK293 cell lines, we hypothesized that these genes might be involved in the editor trafficking process rather than in DNA repair (See Discussion). To demonstrate the role of these genes in increasing editing efficiency, we delivered Cas9-GFP protein into HEK293 cells via lipofection and performed cell staining 6 hours post-delivery of the editor protein (**Figure 4A**, See Methods). Multispectral imaging flow cytometric analysis of the HEK293 cells, knocked out for *BET1L*, *MS4A13*, and *GJB2* genes, revealed an increase in nuclear localization of the Cas9-GFP protein delivered via lipofectamine. Significantly higher nuclear localization of the editor protein, analyzed as a correlation in the fluorescent signals between the nuclear stain and the GFP expression, was observed in the knocked-out lines compared to the WT-HEK293 cells (**Figure 4B, Supplementary Figure S3**). We observed a ∼1.5-fold increase in nuclear co-localization of GFP and nuclear stain signals in HEK293 cells with *BET1L* and *MS4A13* gene knockouts (**Figure 4C**, **Supplementary Figure S4**), suggesting increased cargo delivery into the nucleus.

**Figure 4:**
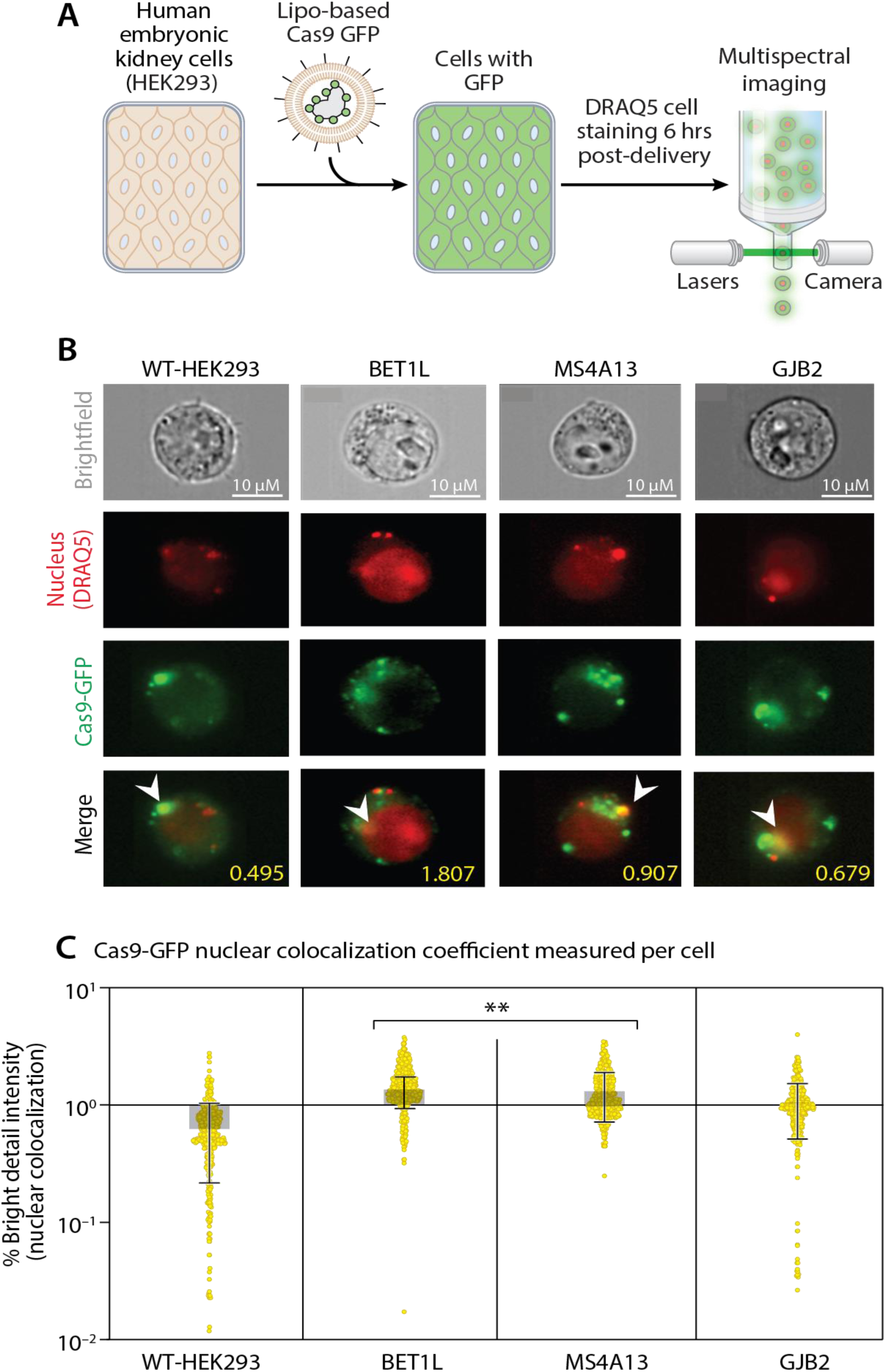
Three gene knockdowns in human cells increase CRISPR-Cas9 editing. See also Figures S3 and S4. **(A)** Schematic workflow of multispectral flow cytometric imaging performed by delivering Cas9-GFP protein in HEK293 cells and KO lines ─ *BET1L, MS4A13,* and *GJB2*. **(B)** Representative single-cell multispectral flow cytometric images of Cas9-GFP transfected HEK293 cells (knocked out for *BET1L, MS4A13,* and *GJB2* genes). The Cas9-GFP here is shown as Green, and the DRAQ5 is shown as Red. Arrowheads indicate the nuclear localization of Cas9-GFP. Numbers in yellow are measured by log Pearson coefficient of overlap between green and red channels. **(C)** Cas9-GFP fluorescent signal localization in the nucleus is defined as the intensity of bright spots in a masked area of an image, which is called as bright detail intensity (BDR). The X-axis shows three different KO HEK293 lines (*BET1L, MS4A13, and GJB2*), WT-HEK293, and UTF conditions. The Y-axis shows a percent increase in the BDR feature in single multispectral flow cytometric images for different conditions. The dot plot shown here represents the nuclear colocalization coefficient (log-transformed) measured per cell for each condition (n=400). Statistical significance was calculated using an ordinary one-way ANOVA Dunnett’s multiple comparisons test (A, B). *, p < 0.05; ****, p < 0.0001; ns, p > 0.05. RNP, Ribonucleoprotein; UT, Untransduced; WT, Wild-type; KO, Knockout; BDR, Bright Detail Intensity; ANOVA, Analysis of Variance.

### *BET1L*, *GJB2,* and *MS4A13* knockdown in iPSC-RPE cells boosts base editing efficiency

To verify the increased editing effect in post-mitotic cells, we used lentivirus to knock out six selected genes in patient-derived iPSC-retinal pigment epithelial (RPE) cells. We confirmed that the RPE cells exhibited >95% transduction (**Supplementary Figure S5A**) corresponding to the gene knockdown in iPSC-RPE cells, and these cells were then edited using lipid-based nanoparticles, targeting the *AAVS1* locus. Subsequent sequencing was performed to analyze the indel rates at the *AAVS1* locus for all transduced wells (knocked out for selected genes) and compared with the untransduced iPSC-RPE cells. Transduced iPSC-RPE cells with BET1L, GJB2, and MS4A13 knockouts showed little or no increase in NHEJ editing compared to control untransduced iPSC-RPE cells. No significant enhancement in CRISPR-Cas9 editing efficiency was observed in iPSC-RPE cells when using LNP-mediated delivery of Cas9 mRNA and sgRNA targeting the AAVS1 locus. Drawing on publicly available single-cell RNA sequencing data for retinal pigment epithelial (RPE) cells, we selected three genes—*BET1L*, *MS4A13*, and *GJB2*—from the six hits identified in validation experiments. The remaining three genes were not expressed in RPE cells, making it clear that they would not influence genome editing efficiency in this specific cell type.

Next, we evaluated the effect of individually knocking down three specific genes on enhancing editing efficiency at a disease-specific locus using adenine base editors (ABE8e)^22,53^. CRISPR base editors modify a single nucleotide in DNA, targeting a precise location in the editing window, without inducing DSBs or requiring the DNA repair machinery. We believed this could provide helpful insight into the functional nature of these genes. We picked the pathological W53X mutation (c.158G>A) in the *KCNJ13* gene, responsible for causing Leber congenital amaurosis type 16 (LCA16) in patients. The W53X nonsense mutation^19,22^ disrupts the Kir7.1 potassium ion channel function in RPE and results in vision loss for LCA16 patients. Here, we investigated the potential of lipid-based nanoparticle-mediated delivery of ABE8e to correct the homozygous *KCNJ13* W53X mutation in gene-knockdown iPSC-RPE^W53X/W53X^ cells. We assessed the extent of phenotype correction achieved with lentiviral-mediated knockdown of *BET1L*, *GJB2*, or *MS4A13* in iPSC-RPE ^W53X^ cells. We performed deep sequencing (**Figure 5A**), immunocytochemistry (ICC), and high-throughput electrophysiology measurements to evaluate the restoration of channel function.

**Figure 5:**
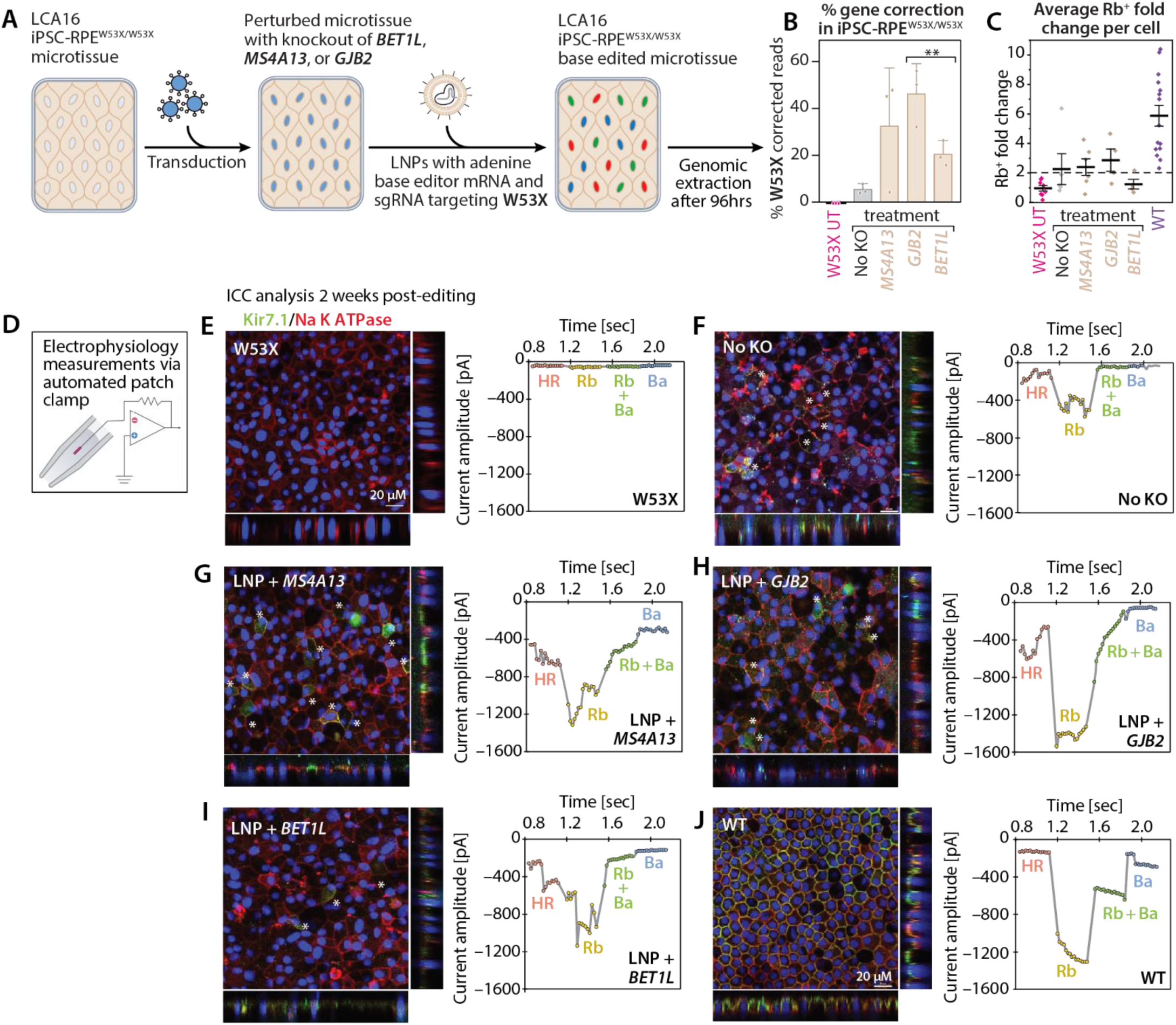
Correction of a pathogenic mutation in post-mitotic, patient-derived RPE is improved by knockout of hits from the CRISPR screen. See also Figure S4. Base editing at the *KCNJ13* W53X locus in a patient-derived Kir7.1 mutant RPE cells via lipid nanoparticles. **(A)** Schematic of base editing, performed by delivering ABE8e base editor to transduced Kir7.1 mutant RPE cells, knocked out for *BET1L*, *MS4A13,* and *GJB2*. **(B)** Data is shown for three replicates (n=3) per treated condition ─*GJB2, MS4A13*, BET1L, untransduced mutant RPE (grey) and untreated/untransduced iPSC-RPE^W53X/W53X^(pink), shown as a correction (TAG->TGG), quantified as the % WT reads out of total reads (Y-axis) for different conditions. **(C)** Average Rb^+^ fold change at −150 mV**. (D)** Schematic of Electrophysiology assay. **(E-J)** Kir7.1 expression (green) and its colocalization with a membrane marker, Na-K-ATPase (red), and a Current-Sweep time plot from a representative cell in different treatment solutions. **(E)** Untreated iPSC-RPE^W53X/W53X^ (n=8). **(F)** Base edited iPSC-RPE^W53X/W53X^ without KO of any gene (n=5). **(G)** Base edited *MS4A13*-KO iPSC-RPE^W53X/W53X^ (n=6). **(H)** Base edited *GJB2*-KO iPSC-RPE^W53X/W53X^ (n=4). **(I)** Base edited *BET1L*-KO iPSC-RPE^W53X/W53X^ (n=4). **(J)** WT iPSC-RPE cells (n=15). Statistical significance was calculated using an ordinary one-way ANOVA Dunnett’s multiple comparisons test (A, B). *, p < 0.05; ****, p < 0.0001; ns, p > 0.05. RPE, retinal pigment epithelial cell; UTF, Untransfected; UT, Untransduced; WT, Wild-type; MT, Mutant; ICC, Immunocytochemistry; ANOVA, Analysis of Variance.

Sequencing analysis in the edited RPE^W53X/W53X^ cells (untransduced) reported 3.9 ± 2% correction of the mutant (TAG > TGG) to wildtype allele, 72 hrs following the LNP-base editing. Surprisingly, the sequencing analysis of edited RPE^W53X/W53X^ cells showed a more than five-fold increase in base editing for the *GJB2* and *MS4A13* transduced conditions (**Figure 5B**), with *GJB2* knockdown increasing the editing to 46.6% ± 5.0%, *MS4A13* knockdown increasing the editing to 32.9% ± 5.0%, and *BET1L* knockdown increasing the editing to 20.9% ± 5.0%. In addition, we performed mock EGFP transduction and utilized the Cas9 genome editor to assess whether the lentivirus-transduction process contributed to an enhancement in editing efficiency. Our results confirmed that increased editing efficiency is directly linked to the knockdown of these selected genes rather than being a result of the transduction process. (**Supplementary Figure S5B-C**).

To investigate the restoration of Kir7.1 channel expression and activity, ICC analysis was done three weeks after the editing using a C-terminal antibody. Our results demonstrated the presence of a full-length Kir7.1 channel (green) in some of the base-edited iPSC-RPE^W53X/W53X^ cells for both transduced and untransduced conditions (treated with base editor). In these cells, the Kir7.1 channel was co-localized with a membrane marker, Na-K-ATPase (red), which confirmed its proper trafficking to apical processes of edited RPE cells, similar to WT cells. The channel was not detected in untreated iPSC-RPE^W53X/W53X^ due to the absence of a C-terminal in the truncated protein product of 52 amino acids.

To assess the functional outcomes of W53X base editing, we performed the automated patch-clamp assay. The cells were exposed to a voltage pulse from −150 mV to +40 mV from a holding potential of −10 mV for 550 ms. Whole-cell current was recorded in different external solutions (physiological Ringer solution; 5 mM K^+^, activator;140 mM Rb^+^ and blocker; 20 mM Ba^2+^) from the pool of transduced and untransduced base edited iPSC-RPE^W53X/W53X^ cells. The current amplitudes at −150 mV were compared to WT and untreated iPSC-RPE^W53X/W53X^ cells. Our reference WT iPSC-RPE cells showed an inward Kir7.1 current of −108.6 ± 17.7 pA (n=15) in 5 mM K^+^ and a 5-fold increased current to - 546.8 ± 87.2 pA in external Rb^+^ solution at −150 mV. On the other hand, the untreated iPSC-RPE^W53X/W53X^ cells, as expected, did not show any activation of Kir7.1 current in the external Rb^+^ solution (Rb fold change 0.96). The pool of base edited iPSC-RPE cells (*MS4A13*-KO, *GJB2*-KO, and *BET1L*-KO) produced a mixed response where some of the cells showed normal functioning K+ channel and an Rb+ fold change above the set threshold, similar to WT, suggesting that these could be perfectly edited (T**A**G>T**G**G) cells without any bystander nucleotide editing. However, in the same pool, some cells appeared mutant with no functional K^+^ channel, as their Rb^+^ fold change was below the set threshold (**Figure 5C and 5D**). A significantly higher (*p-value* 0.009) Rb^+^ fold change was observed in what appear to be perfectly edited *MS4A13*-KO cells (3.5-fold, n=3) compared to our reference untreated iPSC-RPE^W53X/W53X^ cells (n=8). An above-threshold Rb^+^ fold change was observed in two GJB2 KO (4-fold) and one *BET1L* KO (2.2-fold) cells. Kir7.1 profile for a representative cell type is shown in different external solutions as a current-sweep time plot (**Figure 5E-J**). This data from the edited cells (**Figure 5F-I**) demonstrated the increased current amplitude in the external Rb^+^ solution, similar to WT iPSC-RPE cells (**Figure 5J**). Further, the addition of Ba^2+^ in the Rb^+^ external solution caused a decrease in the Rb^+^ signal, which was further decreased in a complete blocker solution of 20 mM Ba^2+^. This response was completely absent in untreated iPSC-RPE^W53X/W53X^ cells (**Figure 5E**).

This data confirms that individual KO of these selected genes does not alter the Kir7.1 property in the edited cells. This further reinforced our sequencing analysis, confirming that the knockdown of these genes significantly enhances editing efficiency in clinically relevant post-mitotic cells without altering the Kir7.1 channel property.

## Discussion

Here, we introduced a large genome-wide CRISPR-based pooled screening approach that links the editing outcome with the guide RNA library on a single ∼150 bp amplicon, enabling us to investigate the factors influencing CRISPR-based editing in human cells. Our novel genome-wide CRISPR screening strategy identified many *host genes* localized to and involved in mechanisms spanning the cell membrane, cytoplasm, several organelles, and nucleus. These mechanisms could generally control cellular uptake, trafficking, and nuclear import of genome editors but can also vary across cell types and identical cells in each tissue or cell population. We could identify genetic dependencies that enhance CRISPR editing efficiency by performing locus-specific deep sequencing of our edited HEK293-Brunello cell population, which had undergone multiple gene perturbations. By comparing two distinct groups—those that received an edit and those that did not—we ranked the genes in our extensive Brunello library, allowing us to assess the effects of nearly every human gene on editing efficiency. We then selected a subset of genes consistently enriched in our MAGeCK analysis, integrating ranked gene function and subcellular localization information. One such hit selected in our analysis was SLCO1C1, responsible for trafficking thyroid hormones in brain tissue.^54^ Knocking out this gene favored the editing process, as did knocking out other genes, such as BET1L, MS4A13, RBM44, GJB2, and *ZNF584*, favored the editing process. All six genes significantly enhanced editing, with a>5-fold increase in editing via LNPs (Cas9 mRNA) and a 2.5-fold increase in editing via lipofection (Cas9 RNP). Our screen identified negative gene hits with significant negative fold-change (FDR ≤ 10%), but genes identified as positive hits did not exhibit significant positive fold change (FDR ≤ 10%). These genes seemed to be essential for NHEJ-based editing in HEK293 cells for lipid-based delivery; for example, the *LIG3* gene was one such missing gene, which encodes for DNA ligase3 and is essential for NHEJ-based DNA repair^55,56^. *POLR1E* is another gene essential for gene editing and encodes a component of RNA polymerase I (Pol I), enabling general transcription initiation factor binding. Other genes listed in the positive gene hits depleted from the edited group involved with DNA repair and damage response were *HDAC2*^60^, *INIP*, and *PSMD3*^57^.

The unique capability of our screen to simultaneously quantify the genetic perturbation identifier (sgRNA) and the editing outcome (phenotype) within a single PCR amplicon makes it particularly suited for studying other next-generation genome editors (such as prime editors and epigenomic editors) and their genetic dependencies to improve editing efficiency and editor delivery. This approach facilitates the characterization of nonviral delivery outcomes for various CRISPR editors within a single pooled experiment. It can be applied to a wide range of editor payloads delivered via nanoparticles, including base and prime editors, in RNPs or mRNA. Understanding the DNA repair mechanisms is crucial for precise genome editing, but the successful "intracellular delivery" of CRISPR components is another significant variable affecting the gene editing strategy. Furthermore, we speculate that knocking out these gene hits might have influenced the cell surfaceome, making knockout cell lines more susceptible to transfections, thus enhancing non-viral delivery. Effective intracellular delivery ensures better uptake of the CRISPR cargo by target cells, significantly enhancing Cas9-mediated editing efficiency. Additionally, the inability of this approach to detect long deletions, translocations, and the loss of other chromosomal segments is one limitation of our genome-wide screen design. Nevertheless, different delivery systems can be tested using our screening design to optimize delivery efficiency in cells from various lineages

Investigating the role of these genes and their contribution to editing efficiency presented a significant challenge. We conducted electroporation studies (**Supplementary Figure S1C**) by delivering RNP cargo to HEK293 cells, including knockout lines for six genes— *BET1L, MS4A13, RBM44, SLCO1C1, GJB2*, and *ZNF584*. Unexpectedly, we did not observe the same trend of enhanced editing in this study. This lack of increased editing efficiency via electroporation indicated that these genes are involved in the trafficking of the editor rather than in DNA repair mechanisms. Further evidence supporting this theory came from base editing studies in iPSC-RPE cells. The base editing nick strategy uses a Cas9 nickase to create a single-strand break at the locus, facilitating the insertion of specific bases without dependence on the cell’s DNA repair machinery. Achieving increased editing via ABE8e editor in iPSC-RPE cells (**Figure 5B**) substantiates our claim that these genes might not be involved in the endogenous repair pathways. This suggests these genes are likely involved in the editor trafficking pathway rather than DNA repair or chromatin arrangement. However, we could only investigate three of the six genes, as the remaining three were not expressed in iPSC-RPE cells. To achieve effective phenotype rescue in post-mitotic cells, such as photoreceptor cells, editing efficiencies of 50% or higher are generally required in the target cells to ensure a meaningful therapeutic impact. By optimizing non-viral delivery methods and targeting these genes using small molecule inhibitors, one can aim to achieve precise genome editing in patient tissues. This presents new opportunities to explore the potential of these genes in other cell types where they are expressed, with the possibility of enhancing editing to correct for other pathological mutations.

## Conclusion

One of the challenges in genome editing is targeting cell types resistant to editing due to their unique repair mechanisms or cellular environments. For instance, post-mitotic cells such as RPE and photoreceptors are notoriously difficult to edit because they lack the active machinery required for efficient DNA repair. The insights gained from the arrayed screen in RPE cells enhancing base editing efficiency^22^ in these difficult-to-edit cell types demonstrated the potential of using the gene targets to develop gene therapy products. Similarly, by identifying key genes and pathways that regulate DNA repair in specific cell types, researchers can develop targeted interventions to improve editing outcomes. For example, temporarily modulating the expression of repair genes in post-mitotic cells could increase the likelihood of successful editing without introducing unwanted mutations. This approach could be particularly valuable in developing therapies for neurodegenerative diseases and other conditions requiring precise editing of post-mitotic cells.

As CRISPR screens continue to evolve, their integration with other emerging technologies, such as machine learning and high-throughput sequencing^61,62^, will further enhance their capabilities and accelerate the development of next-generation genome editors. These advances will transform the field of genome editing, offering new solutions for treating genetic diseases and expanding our understanding of DNA repair and delivery techniques.

## Materials and Methods

### Cell culture

Human embryonic kidney (293T) cells were obtained from ATCC and maintained between passages 15–60 in a growth medium comprising DMEM (Life Technologies), 10% FBS (WiCell), 2mM L-Glutamine (Life Technologies), and 50 U/mL penicillin-streptomycin (Life Technologies). Cells were passaged at a 1:40 ratio using Trypsin-EDTA (Life Technologies) onto gelatin-A (Sigma)-coated plates. HEK293-GFP lines were bought (GenTarget Inc., SKU: SC001), and FACS was performed to isolate cells with a single copy of EGFP, following a previous report^63^. All cells were maintained at 37 °C with 5% CO2 and verified to be mycoplasma-free at least once a month.

Retinal pigment cells derived from human induced pluripotent stem cells (iPSC-RPE) were differentiated as described previously.^64,65^ Briefly, iPSC lines were maintained on Cultrex or Matrigel using mTeSR1 Plus before differentiation. On Day 0 of the differentiation to RPE, iPSCs were lifted using ReLeSR reagent to form aggregates referred to as embryoid bodies (EBs). The suspension cultures of EBs in mTeSR Plus were gradually transitioned to Neural Induction Medium (NIM; 500 mL DMEM/F12 (1:1), 1% N2 supplement, 1% MEM non-essential amino acids, 1% L-glutamine, 2 mg/mL heparin) over 4 days. On Day 7, EBs were plated on Laminin (Cat# 23017015) coated 6-well culture plates, with regular NIM changes until Day 16 when the 3D neurospheres were mechanically lifted off and the remaining adherent cells were allowed to further differentiate in the retinal differentiation media (RDM; DMEM:F12 (3.5:1.5), 2% B27 without retinoic acid, 1% antibiotic-antimycotic solution). The first four media changes from day 16 onwards included 10 µM SU5402 and 3 µM CHIR99021. Between day 75-85 of differentiation, the pigmented RPE cells were purified using Magnetic Activated Cell Sorting as described by Sharma et al.^66^, and plated on laminin-coated 96 well plates or transwell inserts for further experiments. Reagent sources were as follows: Heparin (Cat# H-3149) and SU5402 (Cat# SML0443-25MG) from Sigma-Aldrich; CHIR99021 (Cat# 4423) from Tocris Bioscience, and ReLeSR was from STEMCELL Technologies. All other differentiation reagents were purchased from ThermoFisher Scientific.

### Brunello vector design

A commercially available Brunello lentiviral library was purchased via (Addgene #73179-LV) for our pooled genome-wide screen. This library targets 19,114 genes using 77,441 sgRNA guides, including 1,000 non-targeting guides. The vector was a LentiCRISPRv2 plasmid (Addgene plasmid #52961), a one-vector format expressing puromycin resistance from a 2A site and SpCas9*Sp*Cas9 under the EF1-alpha promoter region. The Brunello lentivirus was designed to target each gene (19,114 in total) using 4 different sgRNAs, with the off-target score and cutting frequency of the library already characterized. The relatively large size of the sgRNA library allowed us to target nearly every human gene and investigate its effect on the editing process through direct perturbation at the Brunello cassette (**Supplementary Figure S1**). By multiplexing next-generation sequencing for this pooled screening approach, we identified sgRNAs that enhance NHEJ at the genomic level, as evidenced by the indel pattern.

### Determination of the Lentiviral Titer

To determine lentiviral titer transductions, HEK293 cells were transduced in 12-well plates with varying volumes of lentivirus (e.g., 0 µL, 30 µL, 60 µL, 90 µL, 120 µL, and 150 µL) with 1.0 x 10^6^ cells per well in the presence of 8 µg/mL polybrene (cat#) in the culture medium. The plates were then incubated at 37 °C. Two days post-transduction, puromycin was added to the culture media at a 1.5 µg/mL concentration in the 12-well plates. Over 10 days, cell viability was counted for different conditions, and a viral dose resulting in 30% transduction efficiency, corresponding to an MOI of ∼0.3, was selected for the large-scale pooled screen (**Figure 1B**). The MOI was calculated for each virus condition as the average cell viability with antibiotic selection divided by the average without antibiotic selection.

### Lentivirus production

Lentivirus for Cas9-mediated editing was manafactured following the protocol outlined by Gándara, Carolina, et al^67^, to knockout the six selected genes in iPSC-RPE cells. In summary, HEK293 cells were seeded and allowed to reach 70% confluence over 24 hours before transfection. At this point, cells were transfected with the target plasmid (containing sgRNA for gene hits, Cas9 protein and puromycin selection marker), **Supplementary Table S1**) and packaging plasmids pMD2.G and psPAX2. The target plasmid carried the target guide RNA for the respective gene hit and Cas9 protein, all regulated by the EF-1α promoter. After transfection, the culture supernatant from HEK293 cells was harvested and frozen at −80 C. The resulting lentivirus titer, determined through functional assays in HEK293 cells, ranged from 10^7 to 10^10^ particles/mL.

### Lentiviral transduction

Transductions in HEK293T cells were performed to represent at least 300 sgRNA per replicate, considering 30% transduction efficiency in ∼100 million cells. To perform the screening experiment, we performed no-spin lentivirus transduction into HEK293 cells at a low multiplicity of infection (MOI ∼0.3). For each well of HEK293 (6-well plate), one million cells were transduced by adding 80 μl of lentivirus (titer =1.3×108 TU) library. Infections were supplemented with 8 μg/mL polybrene and conducted in multiple wells across 6-well plates for ∼100 million cells. After transduction, cells were pooled and resuspended in fresh, complete DMEM media at approximately 0.5 million cells per mL. Cells were then grown for 48 hrs, and subjected to 2 μg/mL puromycin added 2-3 days post-transduction. Cells were selected with puromycin for 48 hrs transduction post-transduction, and media were changed as needed. Dead cells were periodically removed by spinning the cultures from the media and washing them with PBS. The HEK293-Brunello screen library consisting of ∼ 200 million cells was further subjected to Lipo-based CRISPR Cas9 editing as described below.

For iPSC-RPE transductions, iPSC-RPE cells cultured as a monolayer on a 96-well plate or transwell inserts were transduced with a mix of 100 µL RDM (Retinal differentiation media) and 200 µL lentiviral prep for each well or transwell insert. On day 2 of transduction, a complete media change was performed, followed by media changes three days per week until transfection was performed using LNPs delivering the intended genome editors. Cells were imaged pre- and post-transduction to confirm no changes to cell morphology due to transduction.

### CRISPR Screening experiments

For the genome-wide Brunello screen, oligonucleotides containing sgRNA targeting sequences (n=4) were synthesized by IDT. Due to the relatively large size of our Brunello library, we tested four sgRNA performance (**Supplementary Table S1**) to target the internalized Brunello cassette before performing the large-scale CRISPR-mediated editing in 200 million cells. Guides were selected based on the editing efficiency (percent indel reads) at the Brunello cassette, and two guides were selected for inducing indels at the target region on the cassette (**Figure 1C**).To perform this genome-wide screening, we edited ∼200 million cells simultaneously using CRISPR Cas9 protein (RNP, Aldevron Cat# 9212-5MG) mixed in ∼ a 1:1 molar ratio with the selected sgRNA, which targeted the Brunello cassette and delivered the cargo using Lipofectamine 2000 (Life Technologies Cat# 11668030). The cells were harvested four days post-transfection to extract genomic DNA from the pool of cells.

For arrayed experiments focused on studying NHEJ in HEK293 KO lines, cells were harvested 48 hrs post-editing via lipid-based methods (lipofection and lipid nanoparticles). The genomic DNA was extracted from all the conditions and sequenced as described below. (See genomic analysis section)

### Next-generation sequencing library preparation

For the pooled screen, the library was prepared from the cells collected at the end of the editing experiment. The genomic DNA was extracted from ∼200 million cells using the Nucleospin Blood XL (Maxi kit, Machery-Nagel, Cat# 740574.50), processing 50 million cells per column simultaneously. The extracted genomic DNA (∼ 1.5 mg) was purified to remove RNA/protein contamination before library preparation. Polymerase chain reaction (PCR) was performed on the genomic DNA, using Q5 High-fidelity 2X master mix (NEB, Cat# M0492S), to amplify the Brunello guides and the cassette’s target region. Custom PCR primers were used to amplify the cassette for a 150 bp PCR product. The PCR reactions were assembled with 60 ng of template into 20μLof total volume. These reactions were run on a thermocycler with the following program setting: 1 cycle of 30 sec at 98 °C; 5 cycle of 10 sec at 98°C, followed by 20 sec of 72°C and 25 sec of 72°C;5 cycle of 10 sec at 98°C, followed by 20 sec of 71°C and 25 sec of 72°C; 20 cycle of 10 sec at 98°C, followed by 20 sec of 69°C and 25 sec of 72°C; 1 cycle of 2 min at 72°C; 4°C hold. Subsequently, the Illumina seq i5 and i7 adapters(p7/p5) were attached via PCR, and the amplified library was purified using SPRI select Reagent (Beckman Coulter, Cat# A63882). We performed a 0.7X reaction as the effective reaction ratio to purify the final ∼150 bp PCR amplicon. The samples were checked for quality control using a Nanodrop Spectrophotometer, Agilent 2100 Bioanalyzer, and gel electrophoresis. The libraries were sequenced on the Illumina NovaSeq 6000 system (paired-end reads for 150bp, using one SP flow cell). The library amplicon sequenced (read 1) in order had the following lengths: 12nt = UMI sequence; 34nt = Forward Primer; 20nt = Brunello sgRNA; 80nt = on-target region; 22nt = Reverse primer. Due to the large experimental pool of cells, our library preparation protocol was somewhat lossy, but we still achieved sufficient amplicon recovery to ensure good sequencing depth, with a median coverage of ∼200 reads per sgRNA in our NGS analysis. (**Figure 2A, Supplementary Figure S1A**).

For validation arrayed screen experiments (Figure 3A), 26 genes were selected (**Supplementary Table S2)**, and monogenic knockout (KO) lines were generated for each gene in HEK293 and HEK293-GFP lines. To establish knockout lines, we transduced LentiCRISPRv2 plasmid expressing *Sp*Cas9 and the sgRNA under the EF1-alpha promoter region in conjunction with puromycin resistance linked via 2A peptide. Cells were seeded in 12-well plates at a density of 300,000 cells. On day 1, 20 μL of transfection media was added to each well with 200 μL culture media. The transfection media contained 1 μL of Lipofectamine 2000(Life Technologies), 19 μL of Opti-MEM (Life Technologies), and 1 μg of plasmid DNA. The cells were subjected to puromycin selection at 1.5 μg/mL concentration supplemented in culture media. Dead cells were removed, and media was changed periodically. After selecting the different KO lines for 6 days. The cells were plated for the arrayed validation screen at a density of 100 k cells in a 12-well plate. For each condition in the screen, there were three replicates (n=3) for 26 genes, including WT-HEK293 cells as positive control and untransfected control as negative control.

For the lipofection-based editing at the NRL gene locus (**Figure 3A**), we delivered *Sp*Cas9 protein with the NRL sgRNA, inducing NHEJ at the cut-site. RNP complexes were formed in 25 μL of Opti-MEM (Life Technologies) containing 500 ng of Cas9 protein and 20 pmol of sgRNA to transfect 100K cells in a 12-well plate. In a separate tube, 25 μL of Opti-MEM was combined with 0.75 μL of Lipofectamine 2000 reagent and allowed to combine for 5 min. Lipofectamine and RNP solutions were mixed by gentle pipetting and placed aside for 20 min. After this incubation, 50 μL of solution was added dropwise into the well. Media was changed 24 h post-transfection, and genomic DNA was collected after 72 hours, as described in the genomic analysis section.

For the LNP-based editing, lipid nanoparticles were freshly prepared as described below and added dropwise to the 12-well plate at a concentration of 50 ng per well in 200 μL culture media. The cells were incubated, and the media was changed 24 hrs post-transfection. Genomic DNA extraction was done after 72 hours, as described in the genomic analysis section.

### Lipid Nanoparticle (LNP) preparation

The GDLP (Guanidinium-rich lipopeptide) was synthesized following a previous report^68^. Lipopeptides (98.5 mol%) and 1,2-dimyristoyl-rac-glycero-3-methoxypolyethylene glycol-2000 (DMG-PEG2000, 1.5 mol%) were mixed into a 30 mg/mL methanol solution. The payloads (base editor mRNA/ sgRNA=2:1, wt/wt) were dissolved in a 10 mM citrate buffer (pH=5.0) at 1 mg/mL. The payload solution was rapidly pipetted mixed into the lipid solution at a volume ratio of 3/1 during vortex. The lipid nanoparticles were then dialyzed against DPBS 1X for 1 hour using dialysis units (molecular weight cut-off at 20 kDa) and used fresh. The nanoparticle was characterized by dynamic light scattering (Malvern Zetasizer Nano ZS90) for its hydrodynamic diameter, polydispersity index, and zeta-potential (**Supplementary Figure S6**).

### Flow cytometry analysis

Cells were washed, resuspended in 200 μL or 75 μL of FACS buffer, and analyzed on the Attune. We used GFP and DAPI filters to define populations and cells were gated by relative size, shape, singlets, viability, and GFP expression. Percent negative cells for GFP expression were measured using the DAPI and GFP filters and analyzed using FlowJo (**Supplementary Figure S2**). Voltages were established by running wild-type HEK293 cells and GFP expressing HEK293 cells, as respective controls.

### Genomic analysis

DNA was isolated from cells using QuickExtract™ DNA Extraction Solution (Biosearch Technologies, QE0905T) following treatment by 0.05% trypsin-EDTA and centrifugation. The DNA extract solution was incubated at 65 °C for 15 min, 68 °C for 15 min, and 98 °C for 10 min. Genomic PCR was performed according to the manufacturer’s instructions for using Q5 High-fidelity 2X master mix for NGS library preparation as described earlier. Products were then purified using AMPure XP magnetic bead purification kit (Beckman Coulter) and quantified using a Nanodrop2000. Primer sequences are listed in (**Supplementary Table S1**). For deep sequencing, samples were pooled and run on an Illumina Miseq and Illumina Miniseq 2 × 150 bp. Libraries were prepared as described above. (See Next-generation sequencing library preparation section)

### Pooled screen analysis

The genomic DNA was prepped for Next-generation sequencing on NovaSeq 6000 instrument (Illumina, SP flow cell x150bp), by amplifying the targeted region of Brunello cassette from the edited population of cells via PCR. A total of 600 million reads were generated from NovaSeq6000 for the 150 bp read length of the targeting region, as shown in **Figure 1C**. The sequencing reads were then processed to separate the edited transcript from non-edited transcript reads, generating two groups for comparison – edited and non-edited. The editing pattern categories were analyzed by running Cas-analyzer^69^ on FASTQ file data output from Novaseq. According to the data analysis, the fastest repair occurred via NHEJ, where nucleotide ‘A’ was inserted downstream from the PAM sequence at a 5 bp (+/—2) position (Figure 1F). MMEJ repair was the second most commonly occurring pattern in the edited cell population, where −1 bp deletion was observed at the cut site. This finding is following Fu et al.^70^, reporting a high occurrence of similar editing patterns (MMEJ and NHEJ) in K562 and U937 cells.

Due to the sample size differences between the ‘edited’ and ‘non-edited’ read groups, we reduced the large non-edited read group into subsamples to match the sample size of the edited read group (**Figure 2A**). These randomly extracted subsamples from the non-edited read group were run and compared with the edited group through multiple iterations (n=100), and genes that consistently showed up as negative hits were selected based on three factors - frequency of occurrence, cellular localization, and significance. For every iteration, the FASTQ files were analyzed via MAGeCK, read counts were generated and normalized for the Brunello guides identified from both groups (edited group and subsampled unedited group). The guide’s normalized count file was further used for MAGeCK-iNC to identify the enrichment and depletion of genes in ‘edited’ vs. non-edited’ groups. The results were filtered for proliferative and mitochondrial genes such as *HOOK2, PHB2*, etc., and 26 genes were selected for further validation in arrayed-format screens.

### Immunocytochemistry

Immunocytochemistry was done as described earlier to assess the Kir7.1 protein expression in the edited iPSC-RPE cells^71^. Untreated W53X and WT iPSC RPE cells were used as references. Kir7.1 protein was detected using a mouse monoclonal primary antibody (Santa Cruz, sc-398810, 1:200). A cell-membrane marker, Sodium potassium ATPase (rabbit monoclonal, Thermo Fisher, ST0533, 1:500) was used to confirm the localization of Kir7.1 in the membrane. Alexa Fluor 594–conjugated donkey anti-rabbit (Invitrogen, A21207, 1:500) and Alexa Fluor 488–conjugated donkey anti-mouse (Invitrogen, A21202, 1:500) secondary antibodies were used. DAPI (Biolegend, 422801, 1:1000) labeled the nucleus. A confocal microscope (Nikon C2 Instruments) was used to capture the z-stack images and confirm the localization of Kir7.1 protein in the membrane of RPE cells.

### Electrophysiology

An automated patch clamp (Q Patch II, Sophion, Denmark) was used to record the whole cell current (Kir7.1 function) from the edited W53X, untreated W53X, and WT iPSC RPE cells. The cells were washed with 0 Na-CMF solution composed of (in mM) 135 N-Methyl-D-glucamine [NMDG]-Cl, 5 KCl, 10 HEPES, 10 glucose, and 2 EDTA-KOH (pH 7.4). Papain enzyme (MP Biomedicals LLC, Cat#100921) solution (2.5 μl/ mL) was prepared in 0 Na-CMF solution containing (in mg/mL) 0.375 Adenosine, 0.3 L-cysteine, 0.25 L-glutathione, and 0.05 Taurine. To dissociate the monolayer of RPE cells into single cells, transwells were incubated in Papain solution for 45 mins in the incubator at 37°C with 5% CO_2_. Cells were washed with 0 Na-CMF solution three times and dispersed in serum-free media containing 25 mM HEPES (SH media). The cells were centrifuged at 90 g for 1 min and resuspended in SH media. The cells were kept on the instrument’s shaker for 20 minutes before the experiment. The current was recorded using single-hole disposable Qplates through individual amplifiers. A pressure protocol was used to achieve cell positioning (−70 mbar), Giga seal (−75 mbar), and whole-cell configuration (5 pulses with −50 mbar increment between the pulses, first pulse of −250 mbar). The whole-cell current was measured in response to voltage-clamp steps from the holding potential (−10mV) to voltages between −150mV and +40mV (Δ=10mV). More than 60% of the cells completed the experiment. The cells in which the stability was compromised during the experiment were judged by the leak current and excluded from the analysis. The extracellular solution contained (in mM): 135 NaCl, 5 KCl, 10 HEPES, 10 glucose, 1.8 CaCl_2,_ and 1 MgCl_2,_ pH adjusted to 7.4 and osmolarity 305 mOsm. The intracellular solution contained (in mM) 30 KCl, 83 K-gluconate, 10 HEPES, 5.5 EGTA, 0.5 CaCl_2_, 4 Mg-ATP, and 0.1 GTP, pH adjusted to 7.2 and osmolarity 280 mOsm. For rubidium’’Ringer’s external solution, NaCl was replaced with RbCl [140 mM] and used as an enhancer of Kir7.1 current. The data was analyzed using Sophion Analyzer v7.0.58.

### Multispectral imaging flow cytometry

To visualize editor delivery in the HEK293T cells, we used commercially available Cas9-GFP (Sigma Aldrich, Cat# CAS9GFPPRO-250 μg), which were plated at a density of 500,000 per well in a 12-well plate on day 0. On day 1, 20 μL of transfection media was added to cells in 200 μL of maintenance media. 20 μLof prepared transfection media contained 19 μL Opti-MEM (Life Technologies), 1 μLof Lipofectamine 2000 (Life Technologies, cat#11668027) and 20 pmol of Cas9-GFP. The cells were treated 6 hrs before doing the multispectral imaging flow cytometry. On the same day, HEK293 cells were extracted, centrifuged, and stained with 1:1000 dilution of Draq5 dye (Invitrogen, cat # 65-0880-92). After staining, cells were centrifuged and resuspended in 50 μL PBS. Fluorescence was detected on ImageStream X Mark II (EMD Millipore) according to the manufacturer’s instructions. Nuclear co-localization was measured by IDEAS software package (Amnis) using the co-localization wizard. (**Supplementary Figure S3**)

### Statistical Analysis

Statistical significance was determined with P values <0.05 using Prism version 8 (GraphPad). Data analysis was performed using either a Mann-Whitney test, ANOVA, or an unpaired t-test, depending on the data distribution and the number of comparison groups.

## Data Availability Statement

The data sets used and/or analyzed during the current study are available from the corresponding author upon reasonable request.

## Acknowledgments

We acknowledge funding from the NIH, 144 AAN3683-MSN267461, from the SCGE phase 2 award (U19NS132296 grant), MIRA, 144AAI9998-MSN242967, , and McPherson Eye Research Institute Chairs and Professorships to support the work in this project. We acknowledge the University of Wisconsin Carbone Cancer Center Flow Cytometry Laboratory (Attune and BD FACS Aria II BSL-2 Cell Sorter) for the use of its facilities and services. We also thank the members of the Saha lab for their helpful input on experimental design and manuscript preparation. Also, we sincerely appreciate Adam Steinberg’s valuable contributions to the graphic editing of the figures. The contents of this article do not necessarily reflect the views or policies of the Department of Health and Human Services, nor does the mention of trade names, commercial products, or organizations imply endorsement by the US Government.

## Author Contributions

SS and AA designed the sgRNA and primer sequences for the screen design to perform the genome-wide screen. SS performed and analyzed the genome-wide screen in HEK293 cells and subsequent MAGeCK analysis on the screen data. KS and SS selected the gene hits for further validation in HEK293 and RPE cells, where SS performed the arrayed screen experiments. SS collected genomic DNA and performed NGS for the sequencing results for all the screen experiments. SS created all the monogenic knockout lines and analyzed all gene editing optimization studies, multispectral flow cytometry, and lentivirus transduction workflows. DS, MFZ and GH assisted with lentivirus transduction of RPE cells and iPSC-RPE plating, arrayed experiment design, and cell culture. MK assisted in ICC studies for the RPE^W53X/W53X^ lines and performed electrophysiology studies using an automated patch clamp for the RPE data in Figure 5. MZ and AX fabricated the LNPs used for editing the HEK293 and RPE cells for the validation studies. SS wrote the manuscript with input from all authors. KS, SG, DG, and BP supervised the research. All authors read and approved the final manuscript.

