## Supplemental_file_SS for "Genome-Wide CRISPR Screening Identifies Cellular Factors Controlling Nonviral Genome Editing Efficiency": Final Supplemental_information_SS.pdf

### Supplementary Figure S1

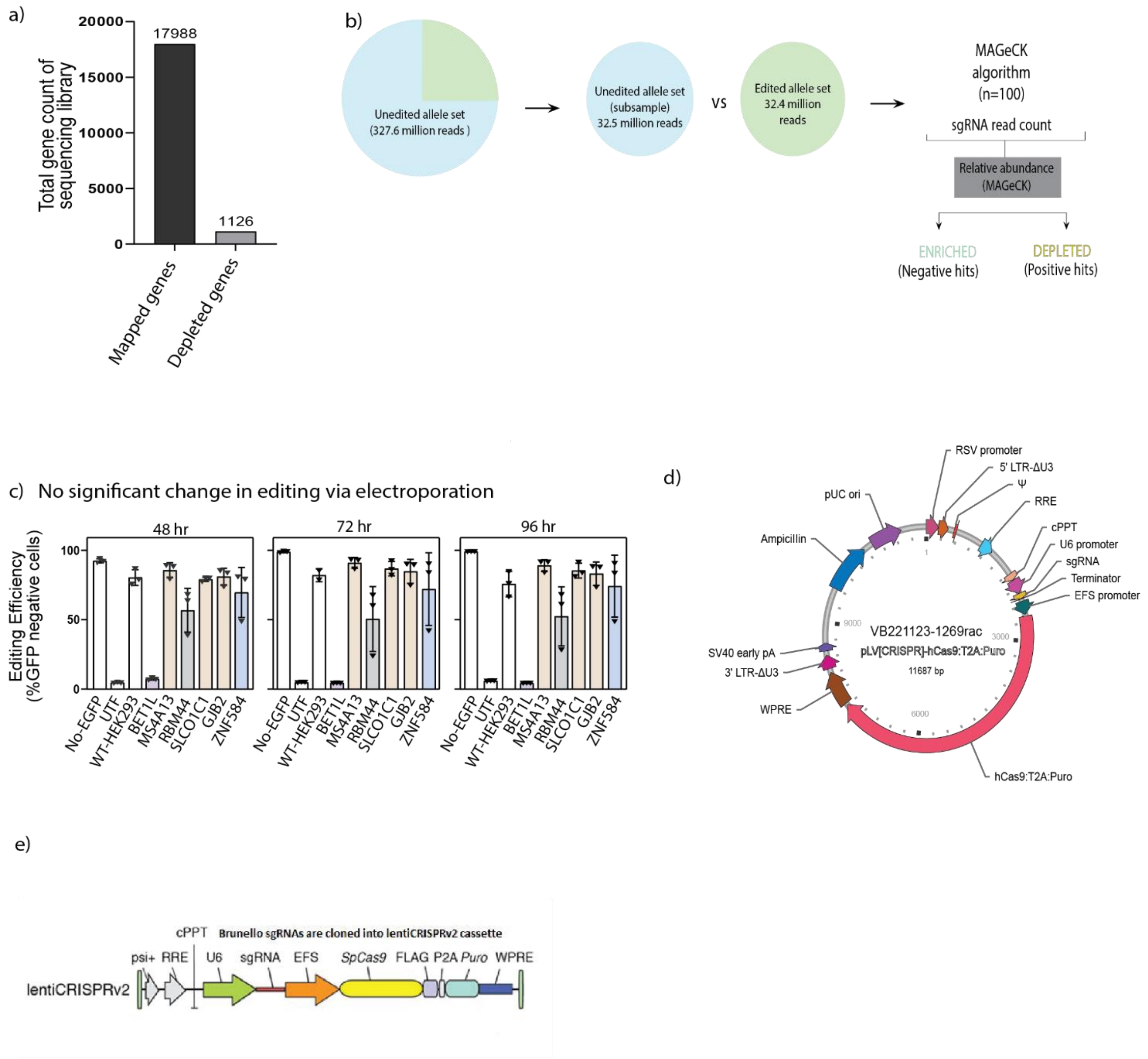


**Figure S1: Screen design and validation of gene hits in HEK293 cells.**
(A) Analysis of the Brunello screen shown here revealed 17988 genes present in the sequencing library, with a median of ~200 reads covered per sgRNA, and 1126 genes were depleted from the total screen library. (B) Schematic of the screen analysis showing the two groups of reads corresponding to the edited reads (NHEJ +) and the unedited reads (NHEJ -). The flow chart shows

the input data for each iteration (n=100) ran through the MAGeCK analysis to shortlist the gene hits. Schematic of the LentiCRISPRv2 plasmid used to create Knockout lines in HEK293 cells and iPSC-derived RPE cells. (C) Electroporation studies shown here are represented as percent negative cells (GFP expression) on the Y-axis for three different time points− 48hr, 72hr, and 96 hr. The X-axis shows the 7 different conditions (n=3) as the different KO EGFP-HEK293 lines (*BET1L*, *MS4A13*, RBM44, SLCO1C1, *GJB2* and ZNF584) with positive (WT- HEK293) and negative control (UTF). (D) Vector builder constructs are used to create monogenic knockout lines for each gene hit for validation studies. (E) Schematic of Brunello internalized cassette.

### Supplementary Figure S2

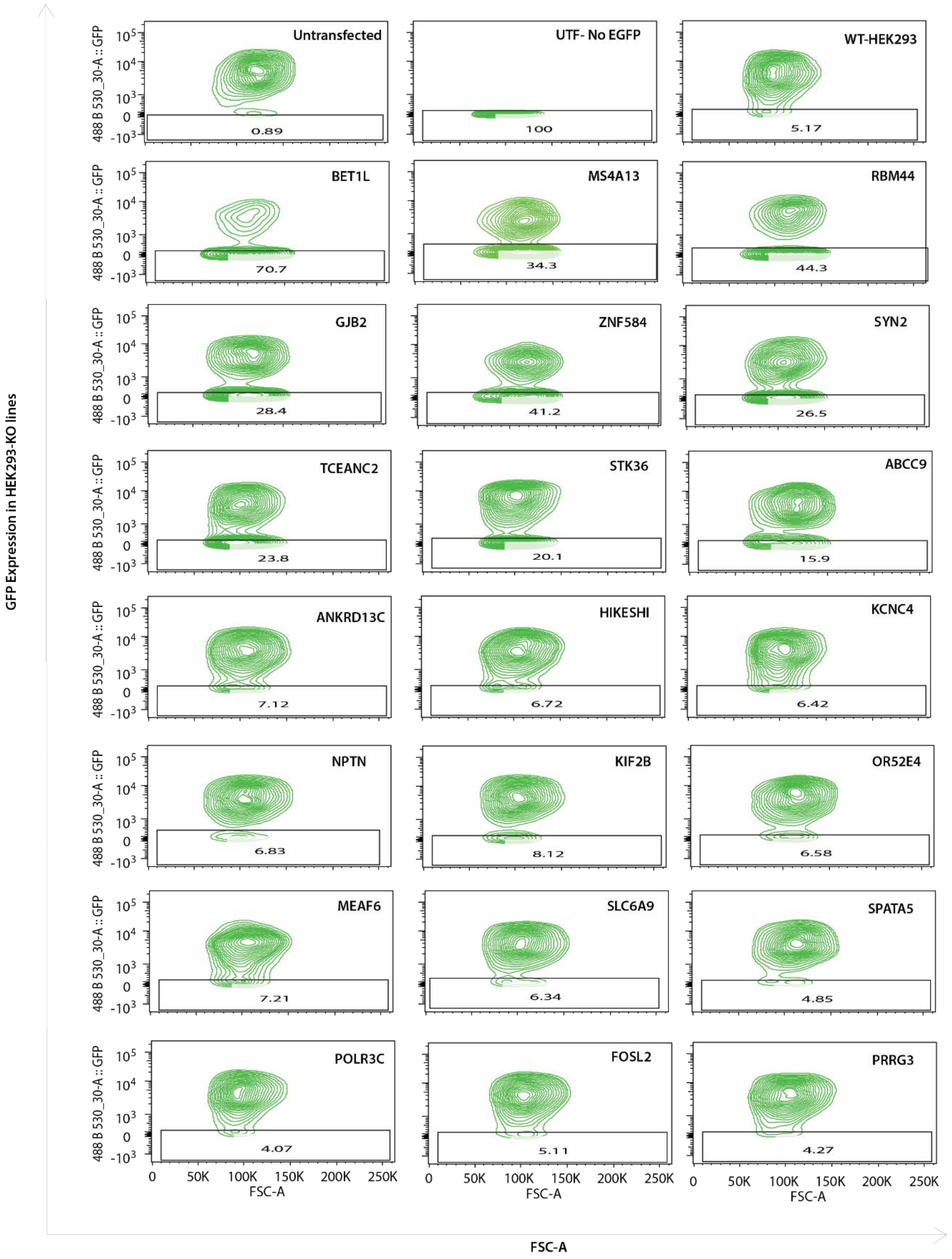

**Figure S2: Flow cytometric analysis done for the KO EGFP-HEK293 via FlowJO.** The flow plots here show the percent GFP negative cells in the boxes for 26 different KO EGFP-HEK293 lines−ABCC9, ANKRD13C, *BET1L*, FOSL2, SYN2, *GJB2*, HIKESHI, KCNC4, KIF2B, POLR3C, PRRG3, RBM44, TCEANC2, OR52E4, NPTN, *MS4A13*, MEAF6, SLC6A9, SPATA5, STK36, ZNF584, UTF (negative control), WT-HEK293 (positive control) and NO-EGFP HEK293 (negative control).

### Supplementary Figure S3

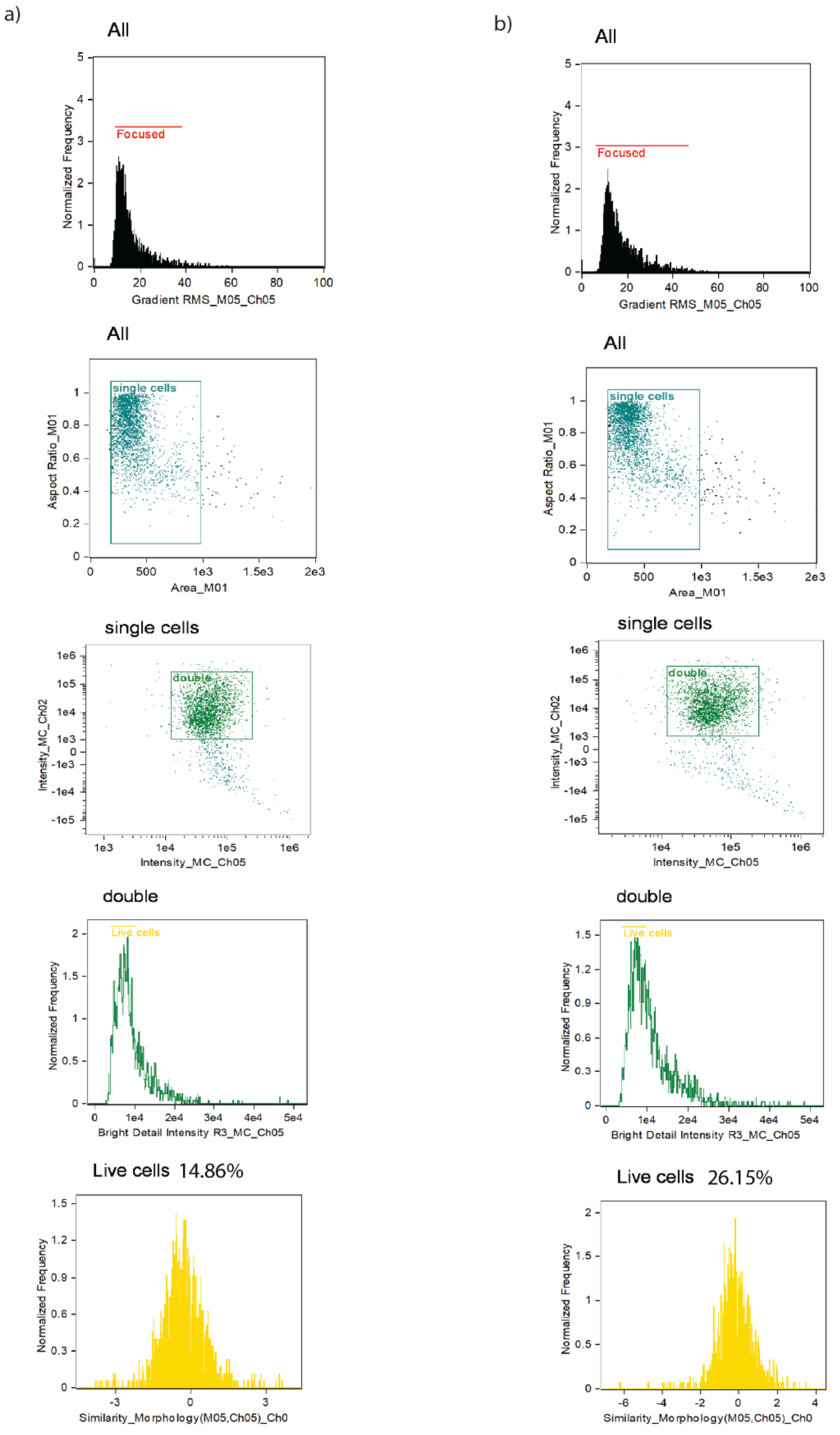


**Figure S3: Gating strategy for multispectral imaging flow cytometry.** Using the IDEAS Software, the gating strategy is employed for the analysis of multispectral imaging flow cytometry to visualize the colocalization of Cas9-GFP (GREEN) and nuclear dye DRAQ5 (RED). Next, the aspect ratio and area measured from brightfield (Ch1) images were plotted against each other to identify single cells. Then, the intensities of Cas9 (Ch2) and DRAQ5 (Ch5) were plotted to identify double-positive cells. Finally, the IDEAS software wizard identified cells with high similarity between Ch2 and Ch5 (Bright Detail Similarity), as shown for (A) WT-HEK293 cells and (B) BET1L-KO HEK293 cells. HEK293 cells were first gated to ensure they were in focus.

### Supplementary Figure S4

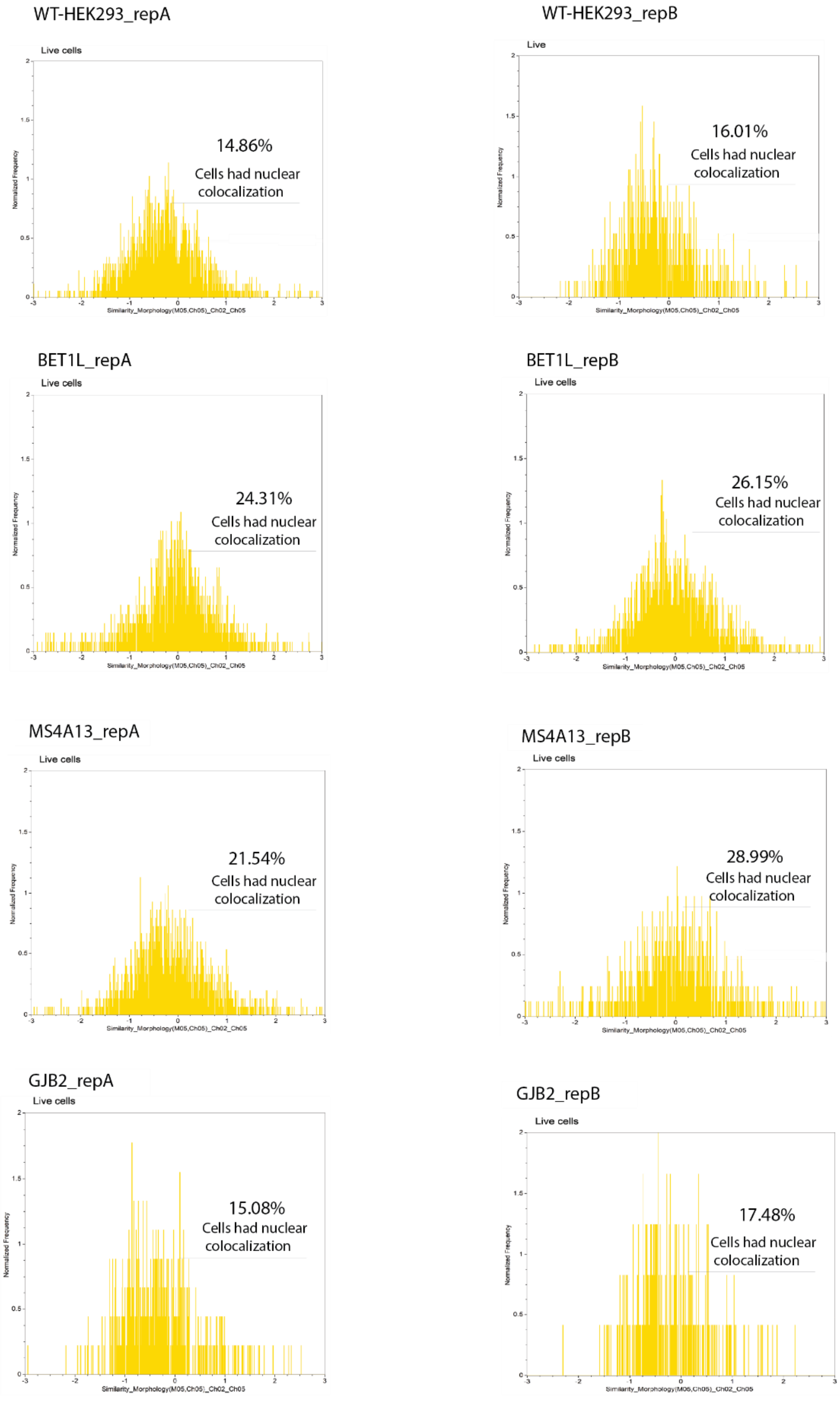

**Figure S4: Nuclear localization of Cas9-GFP protein within HEK cells with knockout of either *BET1L*, *MS4A13* or *GJB2*.** Using gated single cells in multispectral imaging flow cytometry, the colocalization of Cas9-GFP (GREEN, Ch2) and nuclear dye DRAQ5 (RED, Ch5) was measured. For double positive cells, the distribution of Bright Detail Similarity indicates differences among samples from (A) WT-HEK293 cells (B) *BET1L*-KO HEK293 cells (C) MS4A13-KO HEK293 cells (D) GJB2-KO HEK293 cells. The number cell events are on the X-axis and nuclear colocalization on Y-Axis (n=2). See also Figure 4.

### Supplementary Figure S5

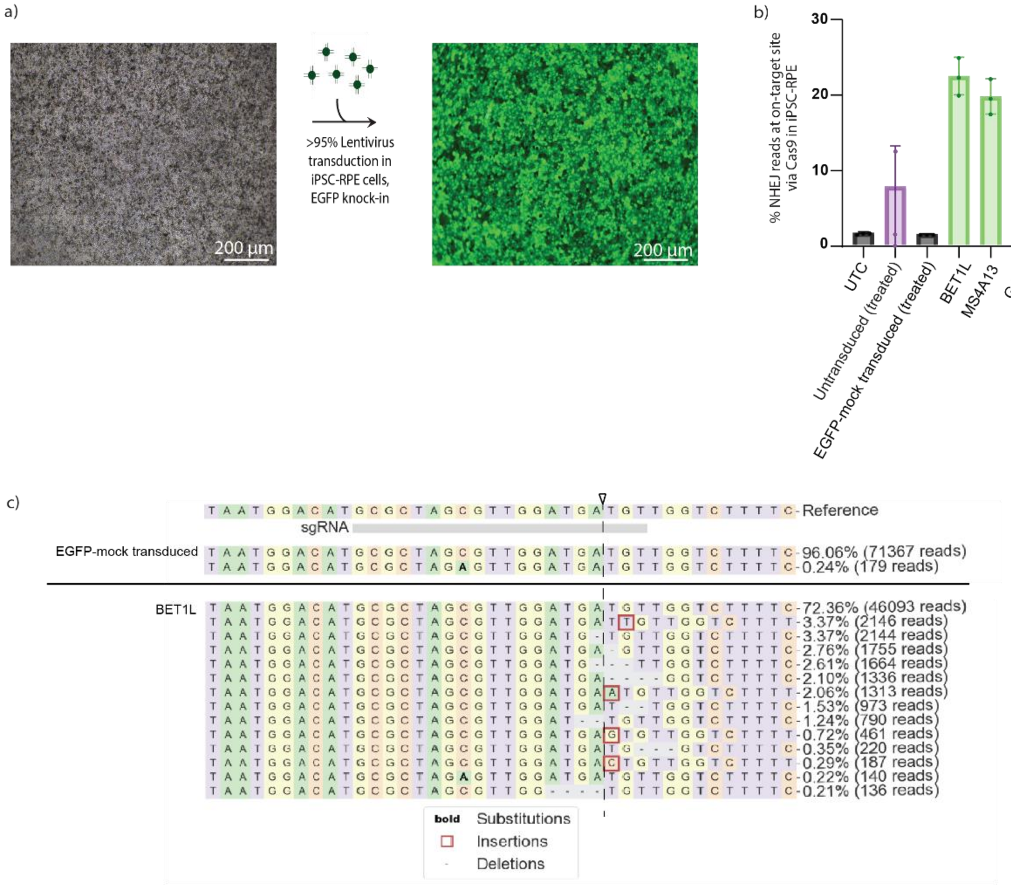


**Figure S5: Base editing at the pathological W53X locus in patient-derived iPSC-RPE cells.** (A) Schematic representing the lentiviral knockdown in iPSC-RPE cells shown as GFP expression.(B) Percent indel frequency generated via Cas9 at the W53X locus in iPSC-RPE cells, comparing 6 conditions- BET1L KO, MS4A13 KO, GJB2 KO, EGFP transduced (Treated), Untransduced (treated) and Untransduced (Untreated) conditions. (C) The indel profile here is representative of the WT corrected reads (TAG ⟶ TGG ) at the W53X locus after treatment with the ABE8e base editor in iPSC-RPE cells (knocked out for *BET1L* gene). Statistical significance was calculated using ordinary one-way ANOVA. Dunnett's multiple comparison test was used as the post-test (C, D, E, and F). *, $p < 0.05$; **, $p < 0.01$; ***, $p < 0.001$; ****, $p < 0.0001$. ANOVA, analysis of variance. RNP, ribonucleoprotein; UTC, untreated and untransduced; KO, Knock-out; NHEJ, Non-homologous end-joining.

### Supplementary Figure S6

**A**

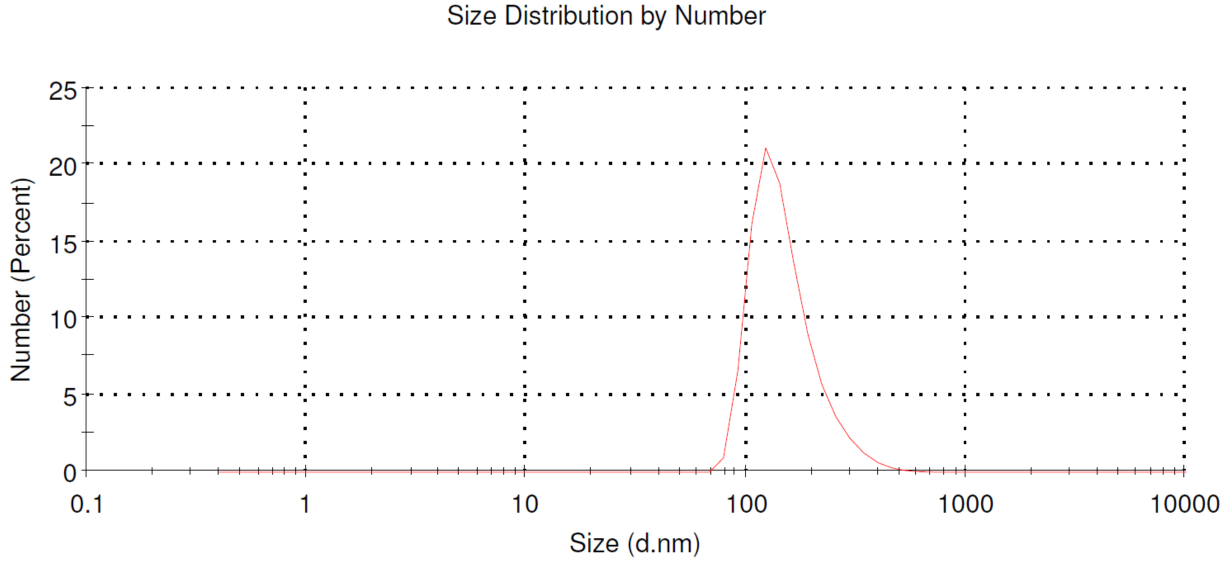


**B**

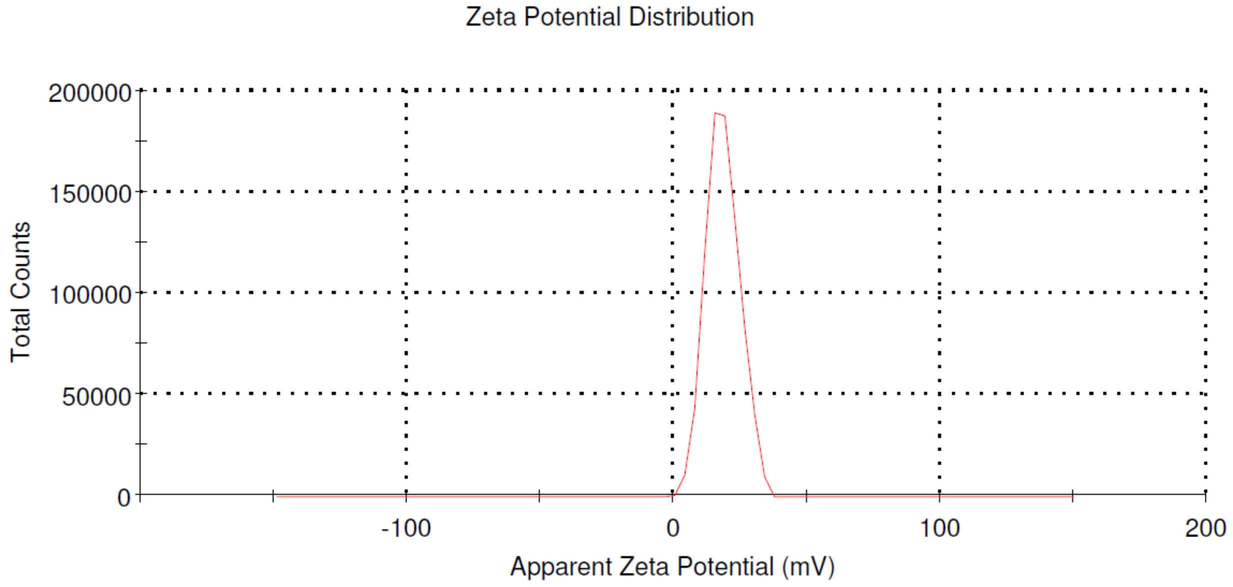


**Figure S6: Size and zetapotential of GDLP LNP.** (A) The mean hydrodynamic diameter of GDLP LNP was 152.7 nm, with a PDI at 0.154. (B) The zeta-potential of GDLP LNP at pH 7.4 was 18.4±6.4 mV.

### Supplementary Table S1

Table S1: Primers and sgRNA used for genome-wide screening

| Construct Name | Sequence (5’ to 3’) |
|---|---|
| Brunello cassette targeting sgRNA 1 | AGAAATAGCAAGTTAAAATA |
| Brunello cassette targeting sgRNA2 | TTATCAACTTGAAAAAGTGG |
| Screen_NGS_Fwd | ATGCATGCTCTTCAACCTCAATAAC |
| Screen_NGS_Rev | CGACTCGGTGCCACTTTTTCAA |
| AAVS1 sgRNA | GGGGCCACTAGGGACAGGAT |
| AAVS1-NGS-Fwd | TGGCTCTGGTTCTGGGTA |
| AAVS1-NGS-Rev | AAGGAGGAGGCCTAAGG |
| EGFP-KO sgRNA | GGGCGAGGAGCTGTTCACCG |
| EGFP-NGS-Fwd | ACAAAAAAGCAGGCTGCC |
| EGFP-NGS-Rev | ACTTGTGGCCGTTTACG |
| NRL sgRNA | GTAAAGCGGGAACCCTCTGA |
| NRL-NGS_Fwd | CAGCCCAGCTCCAGAATGG |
| NRL-NGS_Rev | CTGTAAGGTGTGGAGCCCAG |
| W53X ABE8e sgRNA | GCGCTGGCGTTAGATGATGT |
| W53X-NGS_Fwd | TCAAATGGATGGCGCTCAAAGA |
| W53X-NGS_Rev | ATACCAGAGCACTGCAAAGACAA |
| *BET1L* sgRNA | AGACCGGGAGAACAAGCGAA |
| *MS4A13* sgRNA | GAACCTGTAACTTACAAAAC |
| RBM44 sgRNA | GAAGATAATCGTGTTAACTC |
| SLCO1C1 sgRNA | GGAGCAGGGTGTGTAATCAT |
| *GJB2* sgRNA | GAGTGAATTTAAGGACATCG |
| ZNF584 sgRNA | GTATGGGTCCTCATGTGCCG |

| | |
|---|---|
| *BET1L*-NGS_Fwd | CCCCTCTCTTTTCTCAGCTCA |
| *BET1L*-NGS_Rev | GAGCCTGGTGACTTTGGAGG |
| *GJB2*-NGS_Fwd | CCGGAGACATGAGAAGAAGAGG |
| *GJB2*-NGS_Rev | CCACAGGGAGCCTTCGATG |
| *MS4A13*-NGS-Fwd | TGGGACAAATTAAAGGAGCCT |
| *MS4A13*-NGS_Rev | CTTCTACTATTGGCATGA |
| RBM44-NGS_Fwd | CGGGTTATGAAGTTAAATGTGC |
| RBM44-NGS_Rev | TGAAGTGACTCTTCTTGTCC |
| SLCO1C1-NGS_Fwd | GTTAGCTACTTTGGAGCCA |
| SLCO1C1-NGS_Rev | CCATGAAGAACTGAGGCATTGC |

### Supplementary Table S2

Table S2: The curated list of 26 genes selected for further validation from the genome-wide screening

| Genes | Subcellular location | Function | Reference |
|---|---|---|---|
| *ABCC9* | Plasma membrane | KCNJ11 forms the channel pore, while ABCC9 is required for activation and regulation. | Babenko et.al. [1] |
| *ANKRD13C* | Endoplasmic Reticulum | Acts as a molecular chaperone for G protein-coupled receptors, regulating their biogenesis and exit from the ER. | Parent et.al. [2] |
| *BET1L* | Golgi apparatus | Vesicle SNARE is required for targeting and fusion of retrograde transport vesicles with the Golgi complex. Required for the integrity of the Golgi complex | Tai et.al. [3] |
| *FOSL2* | Nucleus | Controls osteoblast survival and activates CEBPB transcription in PGE2-activated osteoblasts | Chang et. al. [4] |
| *SYN2* | Plasma Membrane | Neuronal phosphoprotein coats synaptic vesicles binds to the cytoskeleton and is believed to function in the regulation of neurotransmitter release. | Thomas Sudhof[5] |
| *GJB2* | Plasma Membrane | Small molecules and ions diffuse from one cell to a neighboring | Choi et.al.[6], Bicego et.al.[7], Oshima et. al.[8] |

| | | | |
|---|---|---|---|
| | | cell via the central pore | |
| *HIKESHI* | Cytosol | It acts as a specific nuclear import carrier for HSP70 proteins following heat-shock stress, which mediates the nucleoporin-dependent translocation of ATP-bound HSP70 proteins into the nucleus. | Kose et.al.[9] |
| *KCNC4* | Plasma Membrane | This protein mediates the voltage-dependent potassium ion permeability of excitable membranes. | Covarrubias et. al.[10] |
| *KIF2B* | Cytosol | Plays a role in chromosome congression | Shrestha et.al.[11] |
| *POLR3C* | Nucleus | DNA-dependent RNA polymerase catalyzes the transcription of DNA into RNA using the four ribonucleoside triphosphates as substrates; Preferentially binds single-stranded DNA (ssDNA) in a sequence-independent manner; Part of POLR3C/RPC3-POLR3F/RPC6-POLR3G/RPC7 heterotrimer coordinates the dynamics of Pol III stalk and clamp modules during the transition from apo to elongation state | Canella et.al.[12] ,Girbig et.al.[13] , Lefèvre et. al.[14] , Wang et.al.[15] |

| | | | |
|---|---|---|---|
| *PRRG3* | Extracellular | Unknown | N/A |
| *RBM44* | Nucleus | Component of intercellular bridges during meiosis. | N/A |
| *TCEANC2* | Nucleus | Unknown | N/A |
| *OR52E4* | Plasma Membrane | Odorant receptor. | N/A |
| *NPTN* | Plasma Membrane | Promotes localization of XKR8 at the cell membrane; Also acts as a chaperone for ATP2B1; stabilizes ATP2B1 and increases its ATPase activity | Suzuki et. al.[16] , Gong et. al.[17] |
| *MS4A13* | Plasma Membrane | It may be involved in signal transduction as a component of a multimeric receptor complex. | N/A |
| *MEAF6* | Nucleus | This modification may both alter nucleosome- - DNA interactions and promote interaction of the modified histones with other proteins, which positively regulate transcription; a Component of the MOZ/MORF complex that has a histone H3 acetyltransferase activity | Ullah et. al.[18] , Doyon et. al.[19] |
| *SLC6A9* | Plasma Membrane | Essential for regulating glycine concentrations at inhibitory glycinergic synapses. | Kim et. al.[20] |
| *SPATA5* | Cytosol | ATP-dependent chaperone, which plays an essential role in the cytoplasmic maturation steps of pre-60S ribosomal | Ni et.al.[21] |

| | | | |
|---|---|---|---|
| | | particles by promoting the release of shuttling protein RSL24D1/RLP24 from the pre-ribosomal particles | |
| *SLC8B1* | Mitochondria | Mitochondrial sodium/calcium antiporter that mediates sodium-dependent calcium efflux from the mitochondrion, by mediating the exchange of 3 sodium ions per 1 calcium ion; Able to transport Ca(2+) in exchange of either Li(+) or Na(+), explaining how Li(+) catalyzes Ca(2+) exchange | Palty et.al.[22] ,Tarasov et. al.[23] , Roy et.al.[24] |
| *STK36* | Cytosol | Controls the activity of the transcriptional regulators GLI1, GLI2, and GLI3 by opposing the effect of SUFU and promoting their nuclear localization; Serine/threonine protein kinase, which plays an important role in the sonic hedgehog (Shh) pathway by regulating the activity of GLI transcription factors | Murone et.al.[25] |
| *ZNF584* | Nucleus | It may be involved in transcriptional regulation. | N/A |
| *MEPE* | Endoplasmic Reticulum | Promotes dental pulp stem cell proliferation and differentiation; Promotes renal phosphate excretion | Rowe et.al.[26] , Martin et.al.[27] |

| | | | |
|---|---|---|---|
| | | and inhibits intestinal phosphate absorption | |
| *SFR1* | Nucleus | Acts as a transcriptional modulator for ESR; Component of the SWI5-SFR1 complex, a complex required for double-strand break repair via homologous recombination | Yuan et.al.[28] , Feng et. al.[29] |
| *SLCO1C1* | Plasma Membrane | Facilitates the transport of thyroid hormones across the blood-brain barrier and into glia and neuronal cells in the brain; Mediates the Na(+)-independent high-affinity transport of organic anions such as the thyroid hormones L-thyroxine (T4), L-thyroxine sulfate (T4S), and 3,3',5'-triiodo-L-thyronine (reverse T3, rT3) at the plasma membrane | Strømme et.al.[30] , Pizzagalli et.al.[31] |
| *FAM13C* | Nucleus | Unkonwn | N/A |
